# Designing High-Affinity Progesterone Binders: Pocket Analysis and Scaffold Selection

**DOI:** 10.64898/2026.02.12.704737

**Authors:** Mohammad Pourhassan-Moghaddam, Bruce A. Cornell, Stella M. Valenzuela

## Abstract

Molecular recognition is a central component that confers detection specificity to all biosensors. The design and use of such molecules require consideration of properties including their affinity and selectivity, plus their ease of production and engineering, for downstream commercial purposes.

Progesterone (P4), is a biomarker that is extensively for various diagnostic purposes. Examples include detection of P4 as an indicator of oestrus in cattle breeding, and ovulation in human IVF programs. P4 is also thought to promote strains of breast cancer, resulting in it being an environmental pollutant of interest.

The present study focusses on *in-silico* molecular docking trials of P4 molecules with proteins such as antibodies and receptors. We describe the geometry of novel P4-binding pockets and predict key residues that favour high affinity and selectivity for P4. The *in-silico* molecular docking trials were performed on various mutants of an anti-P4 antibody that had lost their P4 specificity but retained selective recognition of steroids with structures closely related to cholesterol. Reverse-docking trials permitted the identification of novel scaffolds with favourable P4 binding properties. Future reports will validate the predictions of these studies through wet lab experiments. A further opportunity for this approach is to incorporate a scaffold functionality to permit binding of the protein or receptor to other molecules or sites within a biosensor electrode. These findings, and future studies, will assist in development of enhanced biosensing platforms with custom-designed P4 binders, aiding commercialisation using in-house developed reagents to meet IP requirements and minimise scaling costs. The steroid biotechnology market, valued at over $10 billion, also benefits from novel steroid binder designs, facilitating real-time steroid biomonitoring platforms for optimising steroid bioprocesses.

## 1. Introduction

Progesterone (P4), also known as pregn-4-ene-3,20-dione, is a small steroid hormone that plays a critical role in the regulation of mammalian reproduction, as well as non-reproductive systems. The level of P4 fluctuates in certain diseases such as cancers and physiological conditions such as post-ovulation^1^, and measurement of its level can be used to monitor and diagnose disease states or other physiological changes. Close monitoring and accurate measurement of the level of P4 is needed for accurate drug dosing in patients administered exogenous P4 ^2^ for endocrine therapy, which also allows for deciphering the relationship between administered P4 levels and disease progression. Some cancers, such as breast, are induced and progressed by high levels of P4 in the body^3^, indicating elevated P4 levels as a critical cancer biomarker. Besides being a diagnostic biomarker, accurate and timely measurement of P4 can serve as an important prognostic biomarker, helping to predict disease progression and response to treatment including the efficacy of drugs designed to lower P4 levels. Specific and high-affinity P4-binding proteins are key reagents for development of any accurate P4 assays which can be vital for tailoring personalised treatment plans, as it allows for the adjustment of therapies based on real-time P4 level data, potentially improving cancer patient outcomes. Most proteins used for P4 detection are of antibody origin, with a number currently available from a variety of companies, at relatively high cost Production for commercial use of these proteins is prohibitive due to the labour intensive processes required for their preparation^4^ and similarly, their commercial use is often barred due to intellectual property ownerships by the producing companies^5^. Furthermore, due to the small size of P4, it is not readily immunogenic, and must be conjugated to a carrier protein for the production of antibodies^6^. Administration of the carrier-conjugated P4 as the antigen might also lead to antibodies expressing a higher affinity for the conjugated P4, rather than its unconjugated/free-unbound form. This is the case for DB3 anti-P4 antibody which has a 0.36 nM affinity for the conjugated P4, or Progesterone-11-α-ol-hemisuccinate, and only 1nM affinity for non-conjugated/free P4^7^. A higher affinity for conjugated/bound P4 can result in the incomplete replacement of bound-P4 by the free-P4, which would undermine the accuracy of determining levels of P4 in samples by use of competitive assays. In some other approaches, P4 levels can be measured using cellular assays, where P4 binds to intracellular P4-binding proteins and exerts some biochemical changes that lead to detectable read-outs^8,9^. Another application of P4-binding proteins would be designing P4-responsive reprogrammable metabolic circuits that are stimulated in response to the extracellular P4 levels and produce metabolites of high importance in host microorganisms ^8,10^.

In addition to diagnostic applications, P4-binding enzymes play central roles in many physiological^11^ and industrial level metabolic processes, in which P4 is either produced or used as a substrate^12^. The efficiency of such P4 biotransformation processes can be improved by using enzymes that bind with higher affinity and selectivity to the P4 molecule. It is clear from these examples, that the ability to effectively design novel P4-binding proteins, creates the possibility to develop accurate competitive immunoassays and biosensor formats for quantifying progesterone (P4) across clinically and industrially relevant matrices (e.g., serum, fermentation broth, dairy). Selective P4 binders are the key determinant of competitive assay accuracy: they define the assay’s working range and limit of detection (LOD), while minimising false signals arising from structurally related steroids and matrix-derived interferents in protein- and lipid-rich samples. In addition, appropriate binder bioengineering (e.g., paratope engineering and panel-based selection) can deliberately tune cross-reactivity, enabling conversion of a P4-specific antibody into a class-selective steroid binder for applications where a calibrated “total steroid” readout is required across diverse fields.

### Analyte binding protein

A binding protein is comprised of both the analyte or ligand binding pocket, along with a remaining scaffold domain. The binding pocket is responsible for the binding affinity and selectivity of the binder to the target analyte, also referred to as ligand. Binding pockets are created by the 3-dimensional arrangement of specific amino acids and their respective side-chains that provide a suitable interactive surface compatible with the incoming ligand molecule. The analyte-pocket interaction occurs via a range of non-covalent bonds that requires shape and charge complementarity between the ligand and the pocket^13^. Determination of the 3D structure of a binding pocket is often considered a critical step in comprehending ligand-receptor interactions. The results obtained from such studies can be used to design alternative binders with different/tailored binding properties^14^. Besides binding kinetics (on-rates, off-rates and affinity), the selectivity of the binding pocket is another critical factor that needs to be taken into account for the purpose of developing successful binder-based biosensing strategies. Selectivity is defined as the ability of the binding pocket to recognise its corresponding target analyte molecule, and where there is little to no interaction with non-relevant analytes. At the molecular level, selectivity can thus be defined as high-affinity binding of a binding pocket to target molecules^15^. From this, one can extrapolate that by increasing the structural similarity between two molecules, where one is the preferred ligand, the probability of cross-binding between the non-target molecule and the binder pocket, would increase^16^.

Another important, yet less often considered part of the binding protein is the non-pocket region, often referred to as the scaffold, i.e. protein backbone. The scaffold is responsible for the overall structure and stability of the protein and maintains the correct orientation of the binding pocket, ensuring that P4 binding is unobstructed.

Fragmenting the scaffold into smaller elements can improve the receptor stability and simplify the structure of the biosensor. Single chain variable fragments (scFvs) are fusion proteins that combine two parts of an antibody possessing at least three folds connected by two linker segments^17^. Optimal functionality requires the careful choice of the location of the folds, as well as the length and sequence of the linkers. The complexity of fusion protein design increases as the number of folds or domains increase^18^. Therefore, one strategy for simplifying the design is to use smaller single-domain/fold proteins. Using single-domain proteins such as scFVs eliminates the need for interchain linkers.

Small-sized scaffolds can be readily produced in high yields^19^ and are easy to engineer. Here we describe a systematic investigation and utilisation of *in silico* approaches to determine rules for good P4-binding pockets and suitable scaffold regions that yield small, stable P4 binding proteins that can be readily engineered into various biosensor designs (graphical abstract).

**Graphical abstract:**
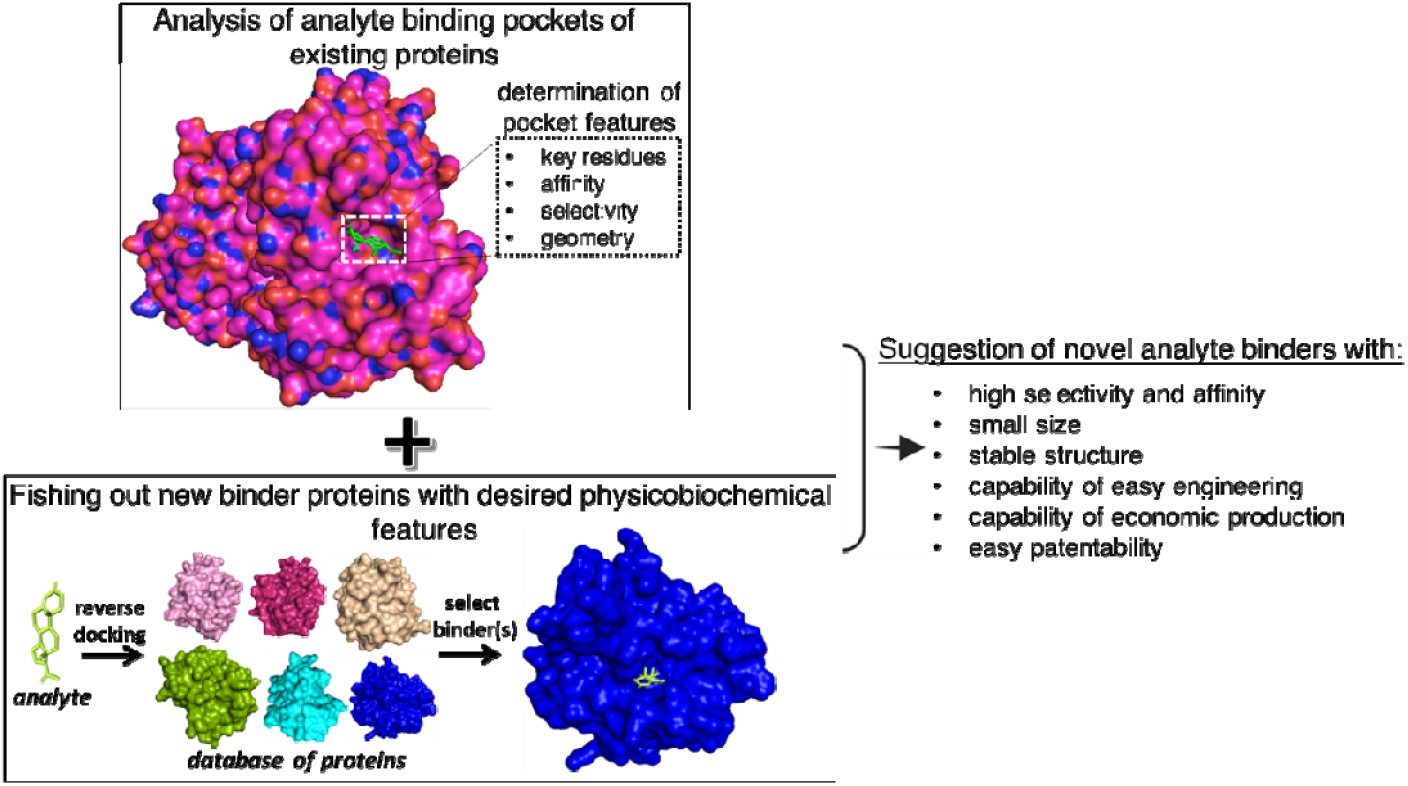
Towards novel engineered P4-binding proteins. Figure generated in PyMol. The design process involves two main steps: (i) characterisation of binding pockets to identify features conferring high affinity and specificity; and (ii) use of these pocket criteria to search for protein scaffolds with superior stability, reduced size, and ease of engineering via reverse docking. The final candidates are proteins that combine the desired binding properties with favourable physicochemical features such as high stability, small size, and cost-effective production potential.

## 2. Methods

### 2.1. Analysis of P4-bound proteins: P4 binding pocket geometry, key amnio acids, and binding properties

The Protein Data Bank (PDB, accessible at: https://www.rcsb.org/) was searched, and 33 P4-bound protein structures were retrieved using the keyword “progesterone” in the search box. The retrieved proteins were imported into Discovery Studio 2020 Client® (DS) and analysed for the interaction of the proteins with progesterone using the (Dassault Systèmes SE, France), tool. The Interacting protein residues, the interaction nature, the specific progesterone chemical groups involved in binding, and the interactions’ distance were all extracted and tabulated. This information was used to suggest a consensus pocket, including binding residues, interaction types, and distances. The shape and biochemical properties of the pockets were also analysed, and proteins were grouped according to their pocket shape and biochemical properties. Furthermore, the volume and hydrophobicity of pockets were calculated using the fPocket tool^20^. Ligand complementarity index between the pocket and the bound ligands was also obtained using Ligand Protein Contact (LPC) tool^21^.

### 2.2. Consideration of P4 binding pocket selectivity for biosensing of total steroids

As P4 shares a considerable similarity with other steroids (figure 1), it is necessaiy to design pockets that are highly P4-selective. Shape complementarity and interaction of pocket residues with the D ring via hydrophobic and hydrogen bonds (C3=0 group)^22^ are critical factors that should be considered while modulating the P4-binding selectivity. For this purpose, the anti-P4 antibody (PDB accession code: 1dbb) was used as a model P4-binding protein to study the binding pocket mutation effect on the binding selectivity of the anti-P4.

**FIGURE 1.**
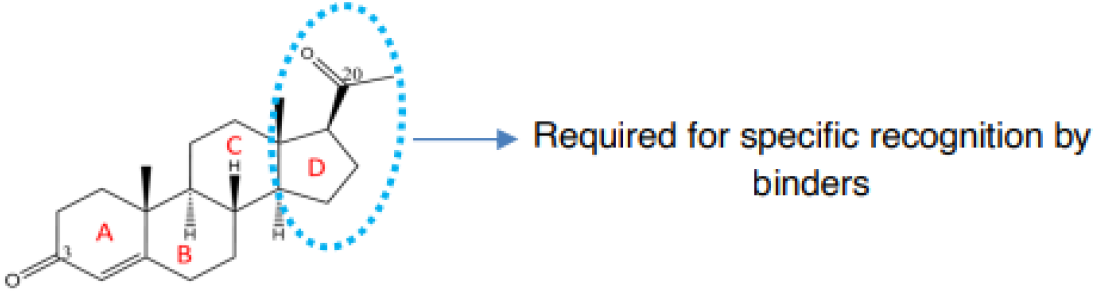
Chemical structure of P4. P4 is composed of four rings and shares structural similarity with other cholesterol-derived steroids. The D ring, however, provides specificity for the P4-binding proteins.

In the first step, computational alanine scanning was used to predict the key residues bound to P4 in the antibody using the ABS-Scan server^23^. The results were analysed according to the difference in free energy of the binding residues between the wild-type complex and the alanine mutants, calculated as ΔAutoDock4.1Score = wild-type Complex_AutoDock4.1Score_ – mutant complex_AutoDock4.1 Score_. An ΔAutoDock4.1Score of 0.039 kcal/mol was calculated from the alanine to alanine mutation score, which was used as the cut-off score/baseline score to exclude the non-important binding residues of the pocket.

Then, critical residues within the binding the pocket were selected based on the alanine scanning output. In the next step, an *in silico* library was created by mutating each critical residue to the other 19 residues. Creation of the library was facilitated by mutateX programme, which provides stability scores for the mutants upon creation^24^. Finally, from the created mutant’s library, only mutants were selected that did not exceed more than 2 kcal/mol in free energies, except for some positions that were kept as control positions-as suggested by mutateX. Next, the structures of the selected mutants were predicted. Swiss Model Server^25^, mutateX outputs^24^, alphafold2^26^ and ABodyBuilder^27^ were used to create protein structures. Energy minimisations were performed by using the default parameters of Rosetta-relax via ROSIE server^28^, and SPDBV tool^29^. Furthermore, optimisations of side chains of structures were performed using FoldX forcefield plugin^30^ integrated with YASARA^31^, and Scwrl4 tool^32^. RMSD values of protein structures were calculated using the align option of the alignment plugin within the PyMOL tool, considering the outlier rejections=5 cycles and cut-off=2. The Align option was selected because the amino acid sequences of the structure were identical for the wild-type models and very close for the mutant models. For the calculation of RMSD values of various ligand binding poses predicted by docking, DockRMSD tool was used^33^. Docking experiments were performed using CB-dock online server^34^ for all six structures of the anti-P4 mutant’s library.

### 2.3. In search of novel P4-binding protein scaffolds with favourable physicochemical properties using reverse docking

A potential small molecule binder should have a relatively deep pocket, similar to the cavity-shaped paratopes found in small molecule-binding antibodies, to accommodate the target small molecules^35^. This feature is also observed in odour-binding proteins^36^ belonging to the lipocalin protein families^37^. Unlike large molecules, small molecules have proportionally smaller surfaces to bind to pockets, and deep pockets can provide maximum contact between the target small analyte and the binder’s pocket. Thus, some of these scaffolds, which are derived from proteins that naturally bind to small molecules, have a greater potential for developing highly selective and high-affinity binders for small molecule analytes. These proteins include lipocalin, single-domain antibodies, odour-binding proteins, and carrier proteins. It is, however, notable that while deep binding pockets favour high-affinity small-molecule recognition, their accessibility can be compromised by tethering or immobilisation linkers required for assay integration, necessitating a balance between binding strength and practical usability. From a scaffold perspective, certain protein frameworks, such as designed ankyrin repeat proteins (DARPins), can be engineered as modular binders due to their repeated structural units^111^. Through engineered multimerisation or surface presentation of DARPin units, multivalent progesterone (P4) binding can be achieved, which is advantageous for signal amplification in immunoassay and biosensor applications. To suggest new scaffolds with potential binding to P4, revers (inverse) docking was used for fishing the potential scaffolds, and then proteins with small size, easy production potential, and high stability were suggested as starting points for developing optimised binders. In reverse docking, a known analyte is simultaneously docked to a library of several proteins to find proteins that can bind to the analyte^38^. The potential of the aforementioned scaffolds for binding to P4 was further tested by undertaking reverse docking, and the most suitable ones were selected.

Reverse docking was performed using P4 as a ligand against the PDB listings as the protein library. PDB was selected as a reference protein structure database to fish out the potential binders because it is the largest depository of the experimentally resolved protein structures, and therefore, the chance for finding suitable scaffolds is maximum. For this purpose, iTARGET^39^ and PharmMapper^40^ tools with default settings were used to perform the reverse docking using P4 as ligand and PDB database as protein library inputs. The output result proteins were screened for suitable scaffold candidates with small size, stable structure, and possibly without cysteine residues.

## 3. Results and Discussion

This study has undertaken by following the steps outlined in the diagram below.

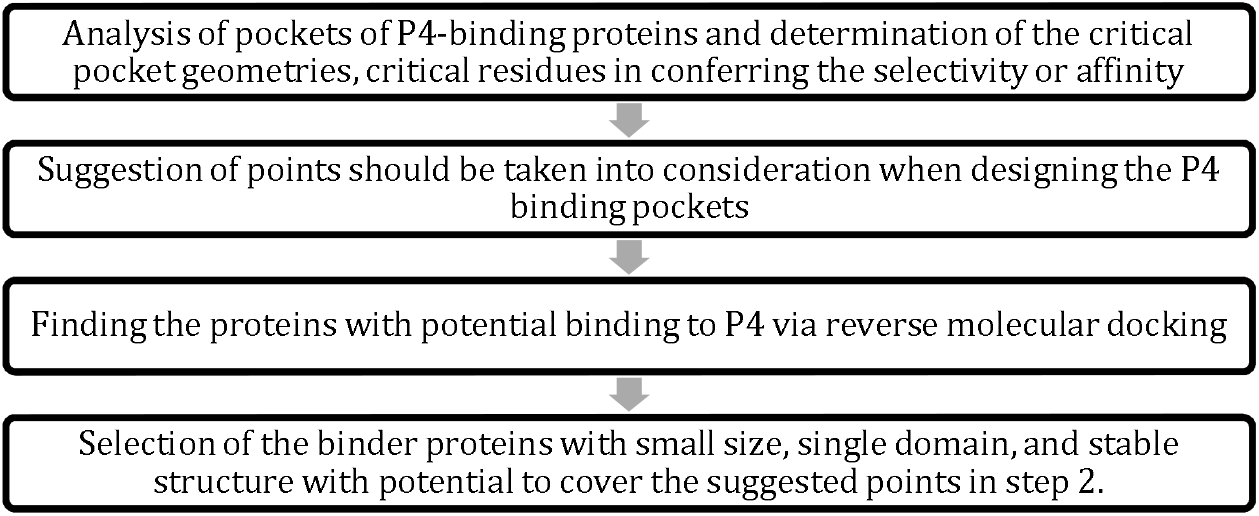

In the first step, published structures of P4-binding proteins taken from the protein data bank, were analysed. The main properties of P4-binding pockets with high specificity and selectivity were extracted from the results. Potential proteins that can bind to P4 were identified using reverse docking, and the potential candidates were selected considering both scaffold properties, including small size, stability and pocket properties obtained from step two.

### 3.1. Analysis of P4-bound proteins: P4 binding pocket geometry, key amnio acids, and binding properties

This section covers P4 binding pocket geometric features critical for selective/high-affinity P4 recognition. Default settings were used for all tools and programs used in this step

The experimentally resolved structures can be used as a starting point to refining P4-binding pockets. Searching the Protein Data Bank (PDB) resulted in the retrieval of 33 protein structures to which P4 was bound. Following importation of the retrieved proteins into Discovery Studio 2020 Client® (DS) tool, the interaction of the proteins with progesterone was analysed. Interacting protein residues, the interaction nature, the specific progesterone chemical groups involved in binding, and the interactions’ distance were all extracted and tabulated (Table 1 below). This information was used to design a consensus pocket, including binding residues, interaction types, and distances. The shape and biochemical properties of the pockets were also analysed, and proteins were grouped according to their pocket shape and biochemical properties. Furthermore, the volume and hydrophobicity of pockets were calculated using the fPocket tool^20^. The volume and hydrophobicity parameters were used to find relationships between selectivity and affinity of the proteins for P4 molecule. Normalised Shape Complementarity index between the pocket and the bound ligands was also obtained using the Ligand Protein Contact (LPC) tool^21^. The resultant Normalised Shape Complementarity index was used to find a correlation between the complementarity of pocket shapes with their affinity and selectivity for their bound P4 molecules. The retrieved proteins belonging to different families, ranging from receptors to enzymes and antibodies, are grouped as shown in Table 1.

**Table 1.** Progesterone-binding proteins retrieved from Protein Data Bank (PDB).

| Protein type | PDB accession code |
| --- | --- |
| Steroid receptors | 1a28, 1ya3, 2aa5, 2aa6, 4fn9, 4ltw |
| Reductase enzymes | 1h60, 1mrq, 2aba, 4l1w, 4l1x, 6gsd, 5mlm |
| Cytochrome P450 | 6f88, 6f8c, 5vbu, 5a1r, 5a1p, 4y8w, 4r21, 1w0f, 4j6c |
| Steroid hydroxylase | 4nkz, 4nky, 4nkx |
| Anti-protease | 6hgk |
| Anti-bodies | 1dbb, 1dbm, 2o5y |
| Apolipoprotein D | 2hzq |
| Engineered avidin | 5lur |
| Steroid binding globulin | 4bb2 |
| Steroid dehydrogenase | 3cot |

Analysis of the P4 binding residues within each pocket revealed leucine (Leu), followed by phenylalanine (Phe), as the most frequent P4-binding amino acids. In contrast, glycine (Gly), serine (Ser) and lysine (Lys) were the least frequent residues. Non-amino acid molecules in the binding site, such as water and Flavin mononucleotide (FMN), also interact with P4 in the binding pocket. Valine (Val) and Leu were the primary nonbinding residues in the binding pocket cavity. Lys, cysteine (Cys) and nicotinamide adenine dinucleotide (NAD) were the least frequent non-binding residues of the analysed pocket cavities. In addition, alkyl and pi-alkyl bonds constitute the primary interactions between the binding amino acids and P4 (figure 2). Figure 3 demonstrates an example of the interactions of progesterone receptor with P4.

**FIGURE 2.**
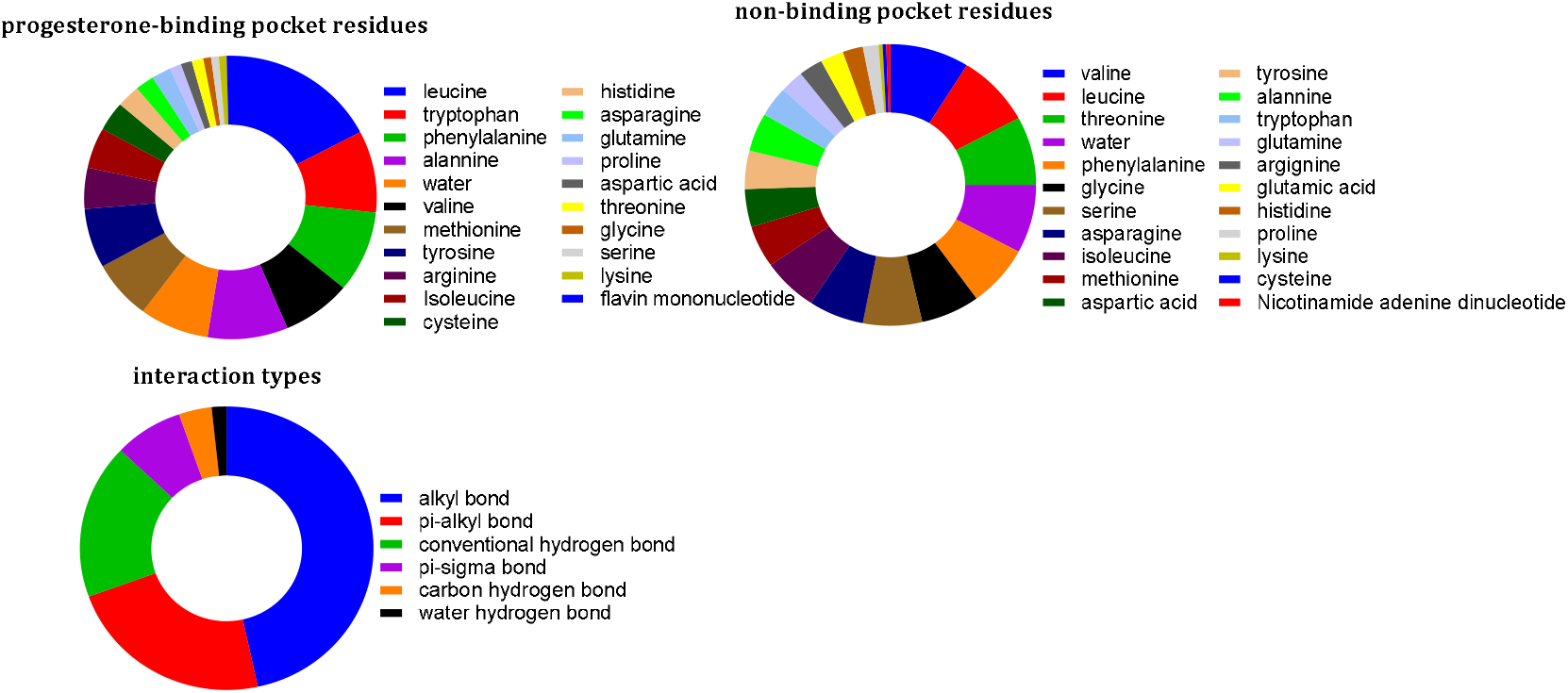
The frequency of P4-binding amino acids, interaction types, and non-binding pocket residues in the P4-binding pockets of the retrieved proteins. Leucine and tryptophan were found among the binding residues as the most frequent amino acids. In contrast, valine and leucine were mostly found as non-binding pocket residues. The hydrophobic alkyl and pi-alkyl bonds were classified as the primary interaction types.

**FIGURE 3.**
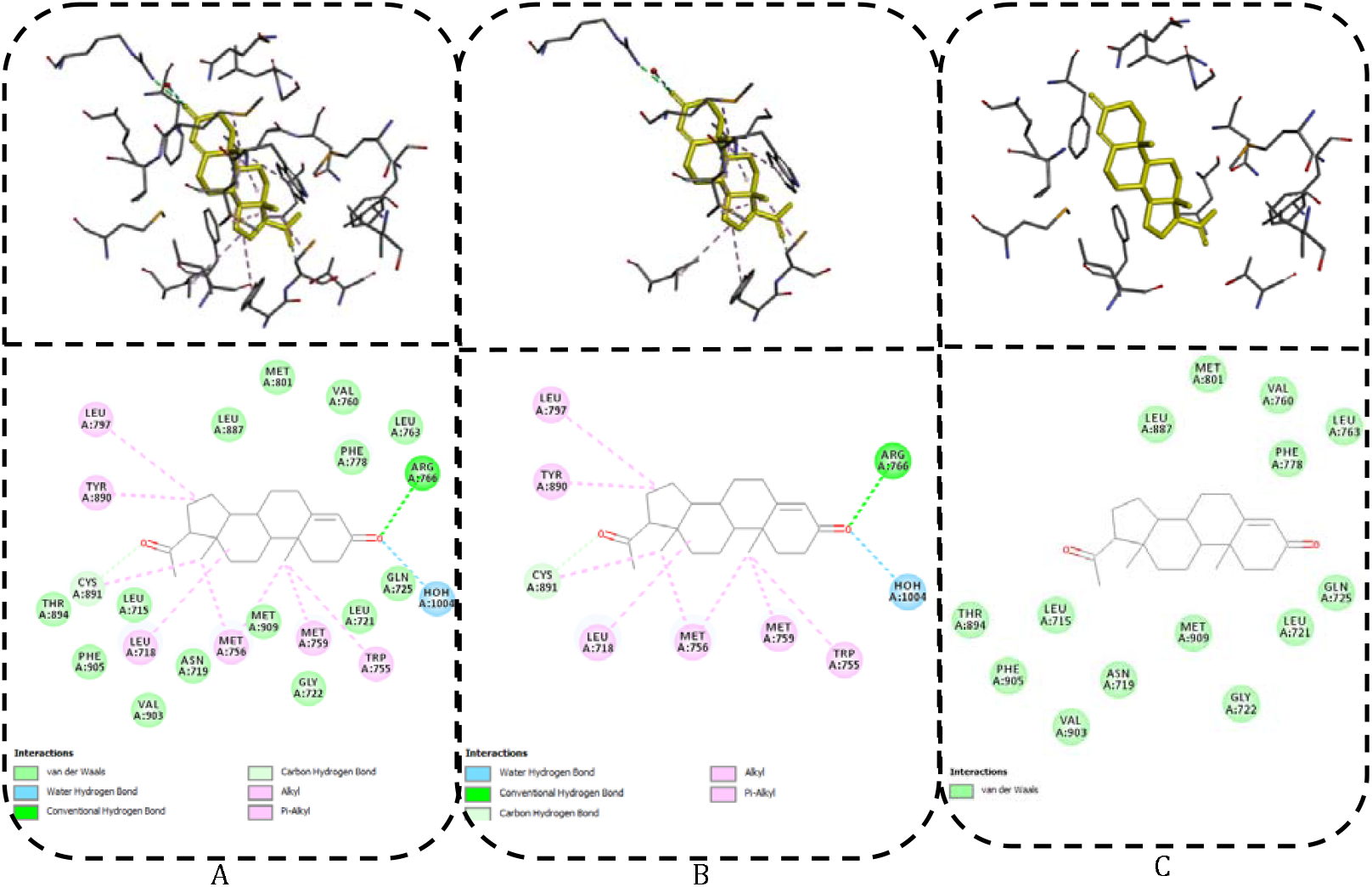
The progesterone-binding pocket of progesterone receptor (1a28) analysed by DS tool. (A): P4-binding and non-binding residues of the pocket. (B): the binding amino acids, (C): non-binding amino acids.

The biochemical properties and interaction geometries of the retrieved proteins are presented in the supplementary tables S1-S18. Interacting amino acid residue and specific P4 site, as well as interaction type and distance, were extracted from the structures of P4-pocket complexes (Supplementary Tables 10A-18A). The information was analysed for commonality, and a consensus P4-binding pocket for the protein groups was suggested (Supplementary Tables 10B-18B). Analysis of the data showed that these pockets contained specific amino acids and interactions. Reviewing these pockets suggested the possibility of improving binding kinetics and reducing protein size via point mutations. For example, analysis of the consensus pocket of anti-P4 antibodies (Supplementary Table. 11B) revealed that almost all consensus residues are located on the variable heavy chain. This might allow for the removal of the variable light chains, hence producing single-domain anti-P4 versions, which are smaller and easier to produce. However, the design of such a single domain structure would require consideration of the non-interacting residues in order to ensure the preservation of the correct conformation of the pocket. Another approach is the inclusion of more hydrophobic residues in order to increase the depth of pocket in the engineered protein, which would increase the contact between P4 and the antibody. Such modifications may increase the affinity of P4 to the engineered anti-P4 antibodies.

Some design strategies could be focused on engineering on- or off-rates, which have implications in designing sensitive and rapid competitive assays. A rapid competition would require the use of P4-binders that possess high on-rates and relatively fast off-rates. In this regard, progesterone receptor would be a good starting candidate (Supplementary Table. 1). However, when P4 binds to the progesterone receptor, the receptor undergoes conformational changes, which results in the burying of the P4 molecule inside the binding pocket^41,42^. Thus, bound P4 requires longer dissociation, resulting in slow-off rates. To increase the off-rate of the progesterone receptor, changing the key residues of the ligand binding domain to prevent the conformational change could be an option. It has been reported that Trp 755 is the key amino acid that is also conserved amongst the nuclear steroid receptors—Trp 755 forms hydrogen bonding with other residues, making the changed structure stable^41^. Hence, mutating this amino acid to glycine, for example, could prevent this kind of stabilisation and might lead to quicker off-rates.

#### A generic pocket for P4

In order to suggest a generic pocket design for P4, which has high selectivity and affinity, the information on consensus pockets was merged (Table 2). According to the pooled data, ten amino acids were present across all pockets with high frequency and provided critical interactions with various parts of the P4 molecule. These amino acids are leucine, tryptophan, phenylalanine, valine, alanine, tyrosine, histidine, asparagine, cystine, and arginine. Five of these ten interacting amino acids were deemed critical and, therefore, would most likely need to be included in the pocket when designing novel progesterone-binding proteins. These five critical residues are leucine, tryptophan, valine, alanine, and tyrosine. Other residues, such as histidine and asparagine, were also considered necessary as important residues in designing new P4-binders and enzymes. Furthermore, the inclusion of histidine in the binding pocket would offer the opportunity to regulate the new synthetic P4-binding proteins with pH^43,44^.

**Table 2.**
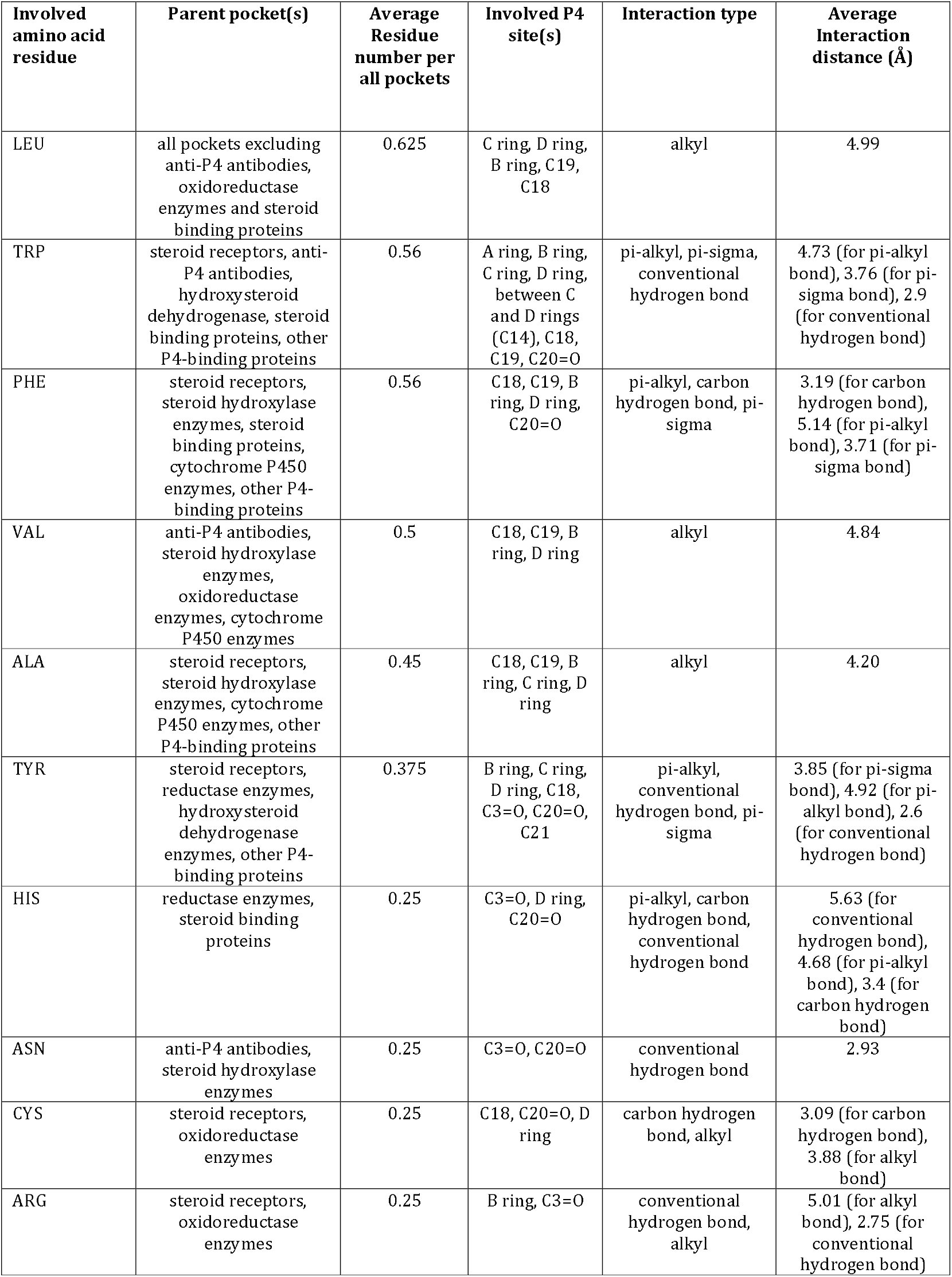
**Common** binding pocket features identified from naturally occurring PG binding proteins.

| Involved amino acid residue | Parent pocket(s) | Average Residue number per all pockets | Involved P4 site(s) | Interaction type | Average Interaction distance (Å) |
| --- | --- | --- | --- | --- | --- |
| LEU | all pockets excluding anti-P4 antibodies, oxidoreductase enzymes and steroid binding proteins | 0.625 | C ring, D ring, B ring, C19, C18 | alkyl | 4.99 |
| TRP | steroid receptors, anti-P4 antibodies, hydroxysteroid dehydrogenase, steroid binding proteins, other P4-binding proteins | 0.56 | A ring, B ring, C ring, D ring, between C and D rings (C14), C18, C19, C20=O | pi-alkyl, pi-sigma, conventional hydrogen bond | 4.73 (for pi-alkyl bond), 3.76 (for pi-sigma bond), 2.9 (for conventional hydrogen bond) |
| PHE | steroid receptors, steroid hydroxylase enzymes, steroid binding proteins, cytochrome P450 enzymes, other P4-binding proteins | 0.56 | C18, C19, B ring, D ring, C20=O | pi-alkyl, carbon hydrogen bond, pi-sigma | 3.19 (for carbon hydrogen bond), 5.14 (for pi-alkyl bond), 3.71 (for pi-sigma bond) |
| VAL | anti-P4 antibodies, steroid hydroxylase enzymes, oxidoreductase enzymes, cytochrome P450 enzymes | 0.5 | C18, C19, B ring, D ring | alkyl | 4.84 |
| ALA | steroid receptors, steroid hydroxylase enzymes, cytochrome P450 enzymes, other P4-binding proteins | 0.45 | C18, C19, B ring, C ring, D ring | alkyl | 4.20 |
| TYR | steroid receptors, reductase enzymes, hydroxysteroid dehydrogenase enzymes, other P4-binding proteins | 0.375 | B ring, C ring, D ring, C18, C3=O, C20=O, C21 | pi-alkyl, conventional hydrogen bond, pi-sigma | 3.85 (for pi-sigma bond), 4.92 (for pi-alkyl bond), 2.6 (for conventional hydrogen bond) |
| HIS | reductase enzymes, steroid binding proteins | 0.25 | C3=O, D ring, C20=O | pi-alkyl, carbon hydrogen bond, conventional hydrogen bond | 5.63 (for conventional hydrogen bond), 4.68 (for pi-alkyl bond), 3.4 (for carbon hydrogen bond) |
| ASN | anti-P4 antibodies, steroid hydroxylase enzymes | 0.25 | C3=O, C20=O | conventional hydrogen bond | 2.93 |
| CYS | steroid receptors, oxidoreductase enzymes | 0.25 | C18, C20=O, D ring | carbon hydrogen bond, alkyl | 3.09 (for carbon hydrogen bond), 3.88 (for alkyl bond) |
| ARG | steroid receptors, oxidoreductase enzymes | 0.25 | B ring, C3=O | conventional hydrogen bond, alkyl | 5.01 (for alkyl bond), 2.75 (for conventional hydrogen bond) |

Other factors that were investigated among P4-bound pockets of the retrieved proteins included volume, hydrophobicity score and shape complementarity between the bound P4 analyte and the corresponding proteins. The obtained results are summarised in table 3.

**Table 3.**
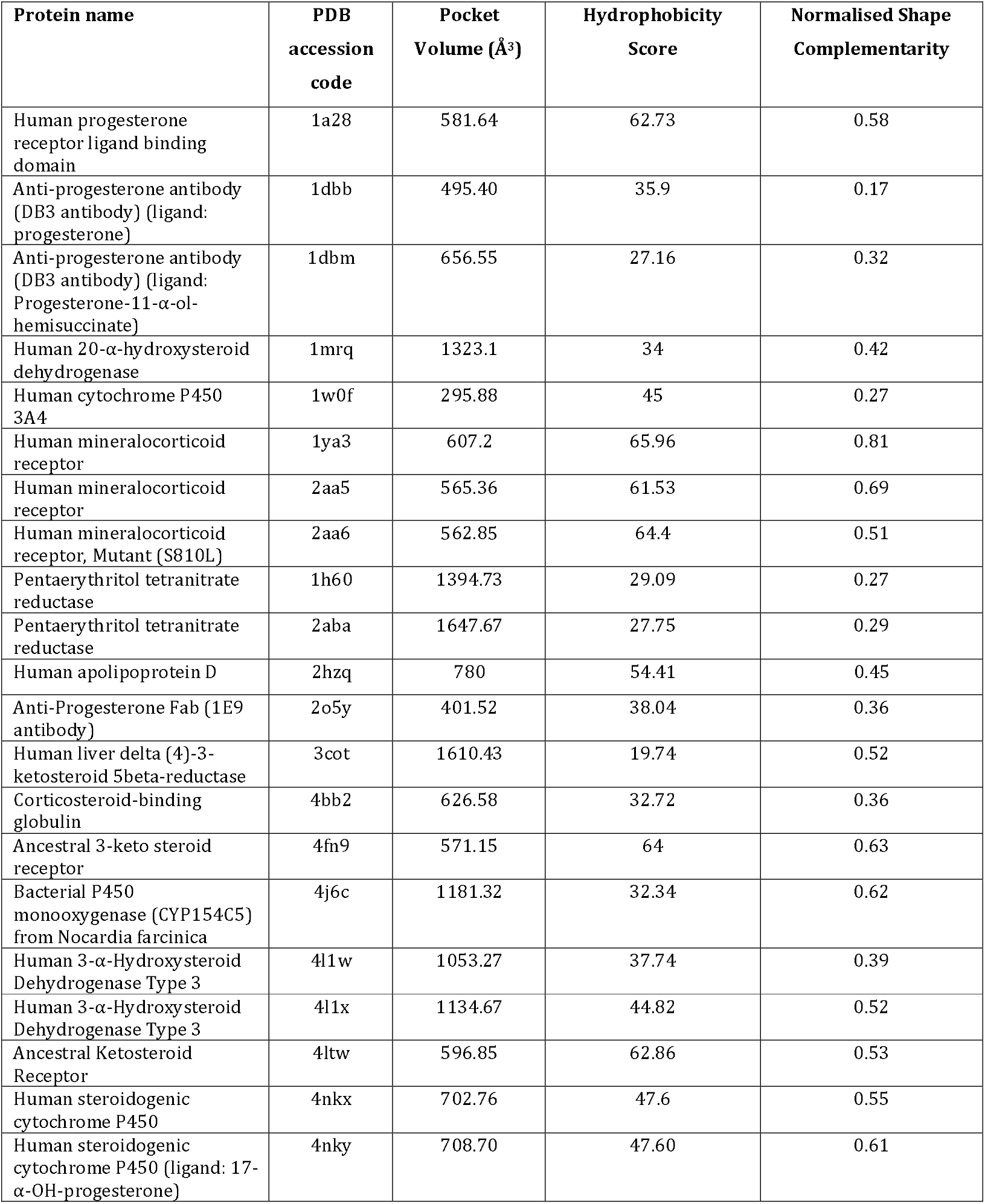
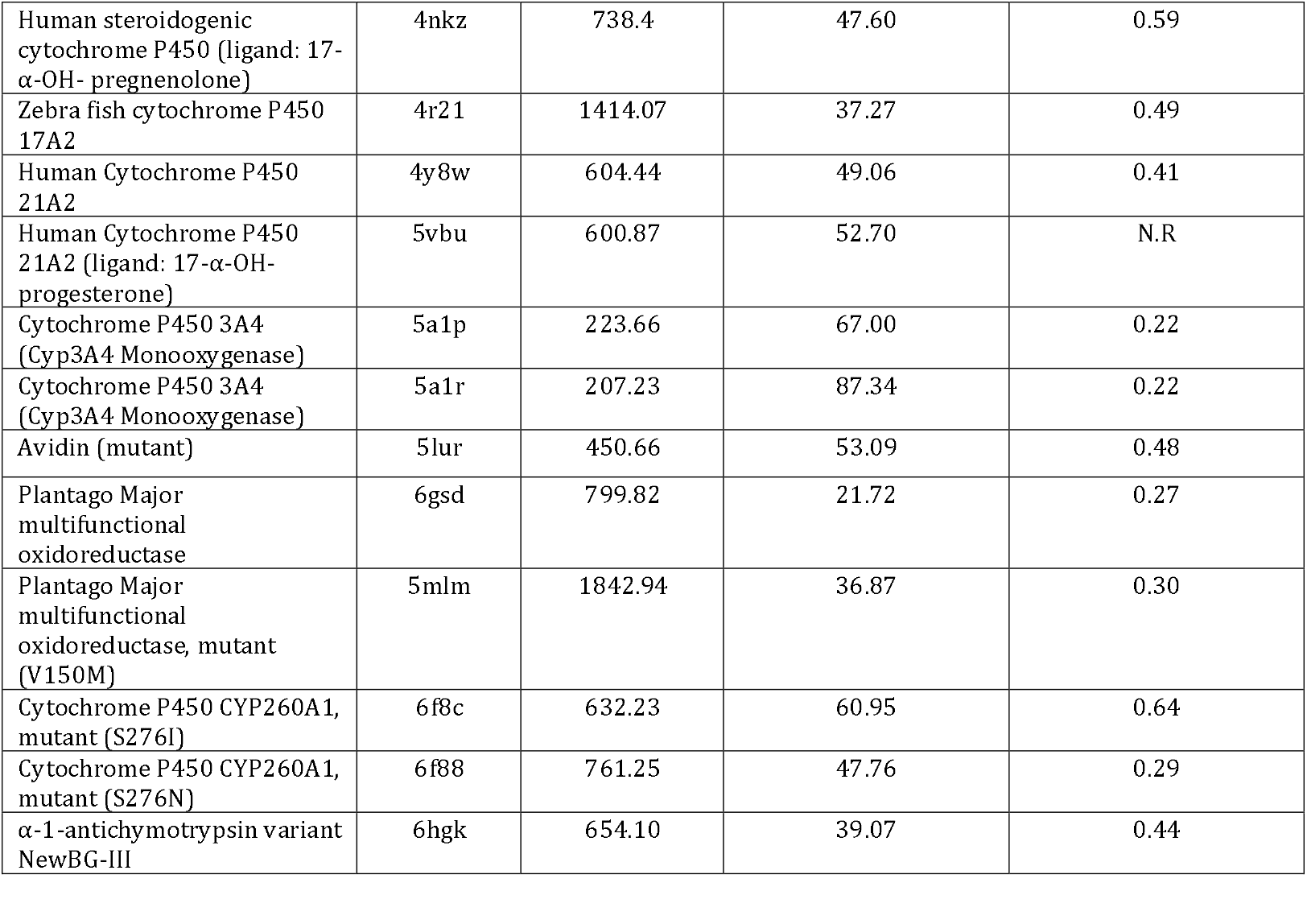
The pocket shape parameters of progesterone-binding proteins.

Table 3 indicates that high-affinity P4-binders such as antibodies and steroid receptors (https://www.rcsb.org/) tend to possess an average pocket size of around 500 Å^3^, around the molecular size of P4. However, it is important to note that other binders may exhibit markedly different binding features because their pocket architecture has evolved for broader specificity or for accommodating ligands with distinct physicochemical properties. For example, some proteins prioritise flexibility or solvent accessibility over strict size complementarity, which can lead to lower affinity for P4 despite having similar cavity volumes.^45^ Another factor is the complementarity between the P4 and the pockets, which are high among the steroid receptors. Pocket hydrophobicity is another factor that is high among the steroid receptors. The two latter factors were found to be relatively smaller for the anti-P4 antibody pockets. Thus, it seems that pocket size is a more important factor in ensuring high affinity binding, which should be considered if alternative P4-binders are to be designed.

In brief, this section focused on the pockets available from the databases, and the information obtained so far can be a preliminary guide for future endeavours in designing steroid-binding proteins. Particularly, common residues involved in pockets of P4-binding (or broadly in steroid-binding pockets) can be a great template when designing or grafting pockets inside other protein scaffolds. Furthermore, the shape and volume of the pockets were found to impact the affinity of binding between protein and P4 molecule.

### 3.2 Consideration of P4 binding pocket selectivity for biosensing of total steroids

#### 3.2.1. Prediction of key P4-interacting residues via computational alanine scanning

Table 4 (the second table in table S19) shows the important residues classified according to the alanine scanning mutagenesis strategy. This alanine scanning score for each protein was obtained by performing in silico alanine scanning to predict the key residues bound to P4The residue with the most negative score was selected as the most important residue of the pocket

**Table 4.**
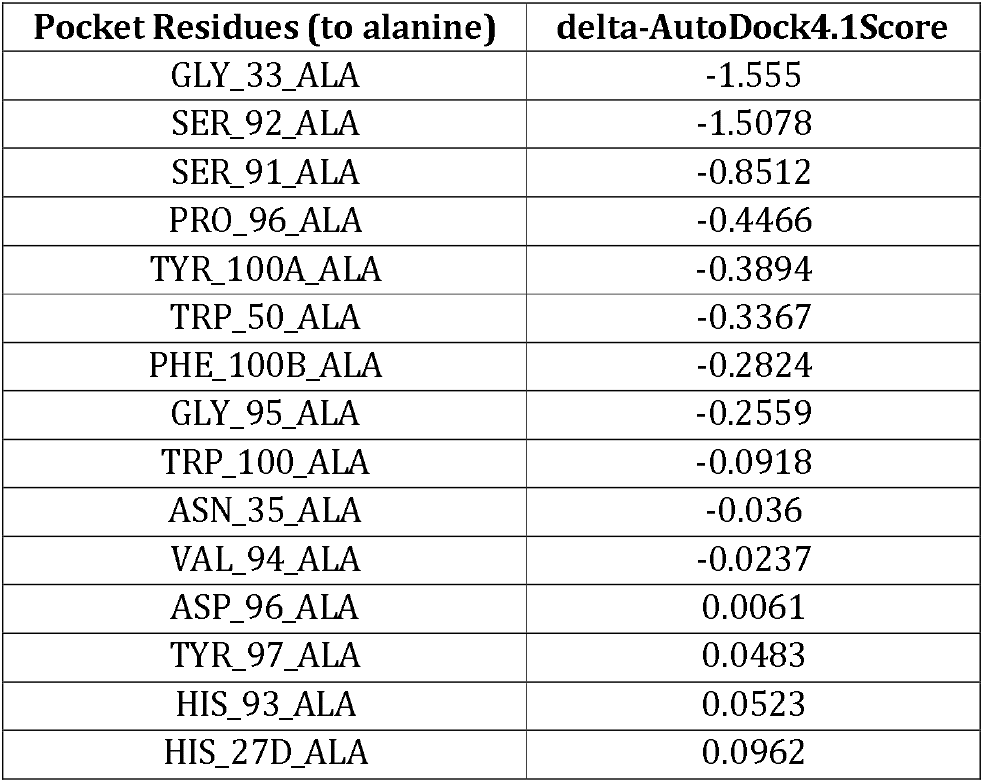
Results of computational alanine scanning of ani-P4 antibody.

#### 3.2.2. Creation of *in-silico* library and docking

In order to further study the results obtained from the alanine scanning, the mutateX programme was employed to produce a library of point mutations based on the mutated anti-progesterone antibody (PDB access code: 1dbb), which served as a model P4-binding protein.

A mutant library was created by considering the results of the alanine scanning of anti-P4 antibody (PDB access codes 1dbb and 1dbm). WH100, WH50 and GH33 residues were selected because alanine scanning showed that mutating these residues might lead to decreased binding affinity. SL91 and HL27 were also selected because alanine scanning results indicated that mutations at these positions might result in higher binding affinity than wild-type antibody. In addition to these residues, FH100 and NH35 residues were selected as they recognise the D-ring of P4, which gives P4 selectivity to the antibody. In the next step, each selected residue was mutated to other residues, provided that the stability of the resultant mutant antibody is not reduced by more than 2 kcal/mol-as suggested by mutateX analysis. The mutateX results suggested GH33 residue as the most unfavourable position to mutate (figure 4). Nevertheless, GH33 was deliberately mutated to other amino acids in order to study the relationship between the stability of the GH33 mutants and their binding to P4.

**FIGURE 4.**
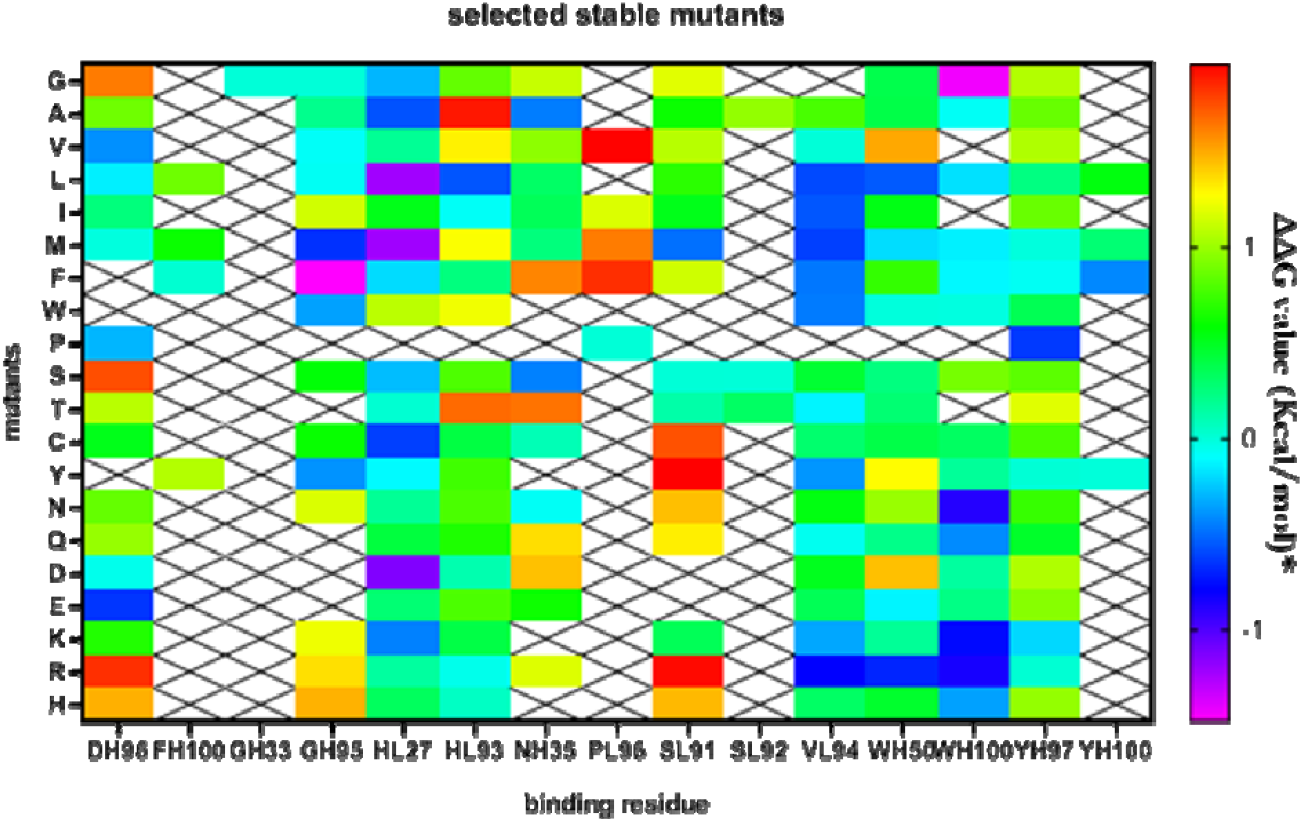
In silico library of stable mutant DB3 anti-progesterone anti-bodies, as suggested by mutateX (*ΔΔG value=ΔG_parent antibody_-ΔG_mutant antibody_).

In the next step, the structures of the mutants and non-mutant (wild type, WT) anti-progesterone antibody were created. A number of methods were benchmarked for structure creation, and it was found that the homology modelling via Swiss-Model server^25^ created the most accurate structures for our purpose, as suggested by excellent RMSD scores and results of trial docking experiments. Furthermore, side chain optimisations did not result in lower RMSD scores (table 5); thus, it was not pursued. As six structures of anti-P4 antibody for an identical sequence are available in the protein data bank (1dba, 1dbb, 1dbj, 1dbm, 1dbk and 2dbl), each mutant and WT structures were modelled using all these structures as templates.

**Table 5.**
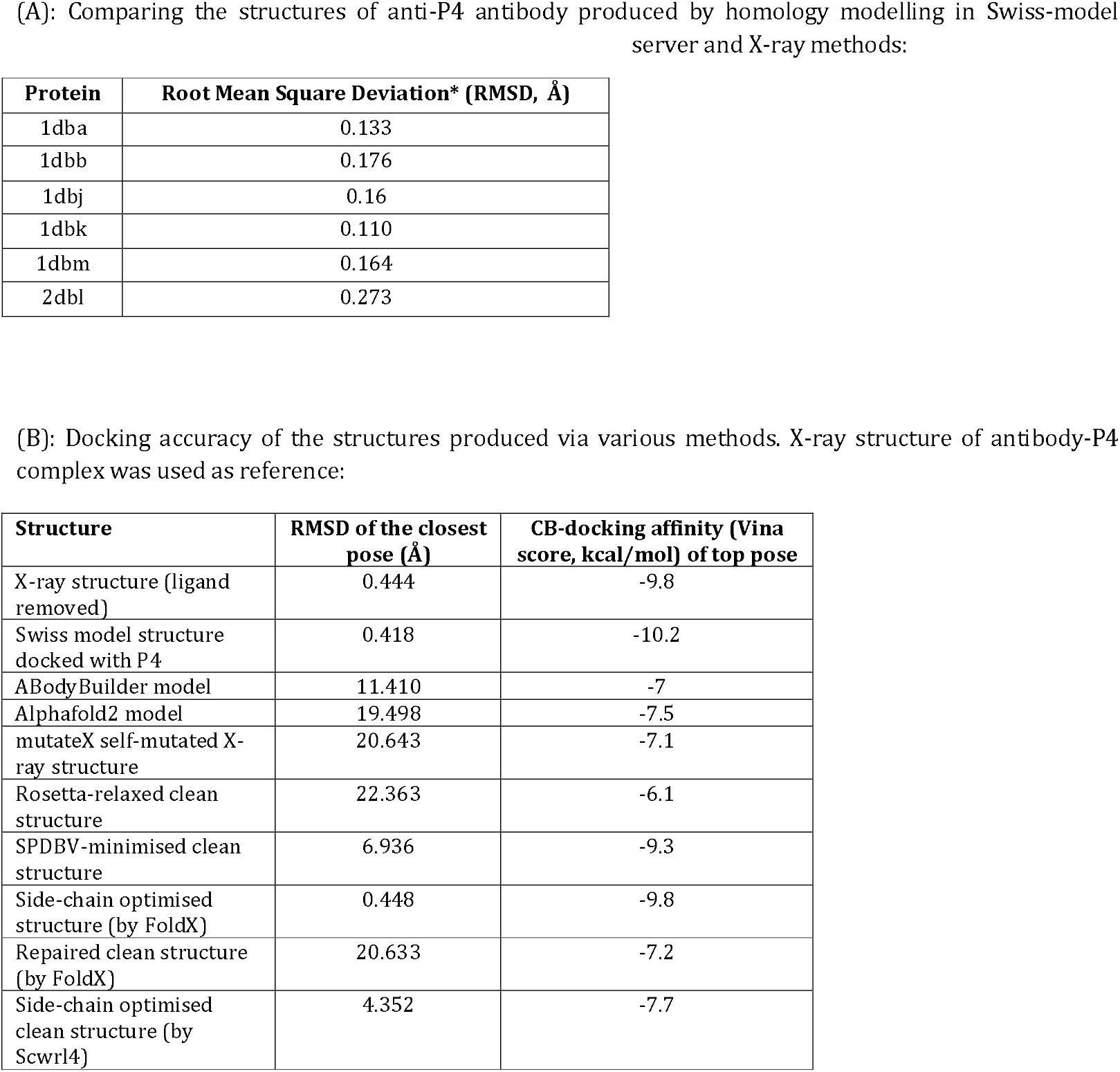
validation of structure prediction method.

| Protein | Root Mean Square Deviation* (RMSD, Å) |
| --- | --- |
| 1dba | 0.133 |
| 1dbb | 0.176 |
| 1dbj | 0.16 |
| 1dbk | 0.110 |
| 1dbm | 0.164 |
| 2dbl | 0.273 |

| Structure | RMSD of the closest pose (Å) | CB-docking affinity (Vina score, kcal/mol) of top pose |
| --- | --- | --- |
| X-ray structure (ligand removed) | 0.444 | -9.8 |
| Swiss model structure docked with P4 | 0.418 | -10.2 |
| ABodyBuilder model | 11.410 | -7 |
| Alphafold2 model | 19.498 | -7.5 |
| mutateX self-mutated X-ray structure | 20.643 | -7.1 |
| Rosetta-relaxed clean structure | 22.363 | -6.1 |
| SPDBV-minimised clean structure | 6.936 | -9.3 |
| Side-chain optimised structure (by FoldX) | 0.448 | -9.8 |
| Repaired clean structure (by FoldX) | 20.633 | -7.2 |
| Side-chain optimised clean structure (by Scwrl4) | 4.352 | -7.7 |

Because an excellent agreement was found between the docking results of the modelled or X-ray structure and the X-ray structure of the antiP4 antibody-P4 complex, CB-dock server^34^ was used to perform all docking experiments on the antibody library comprised of 738 modelled structures. After analysis of the docking scores, no mutation with significantly lower scores compared to the WT proteins was found. However, statistically, increased docking scores suggested that some mutations might lead to decreased affinity or selectivity of DB3 anti-progesterone antibody (figures 5 and 6).

**FIGURE 5.**
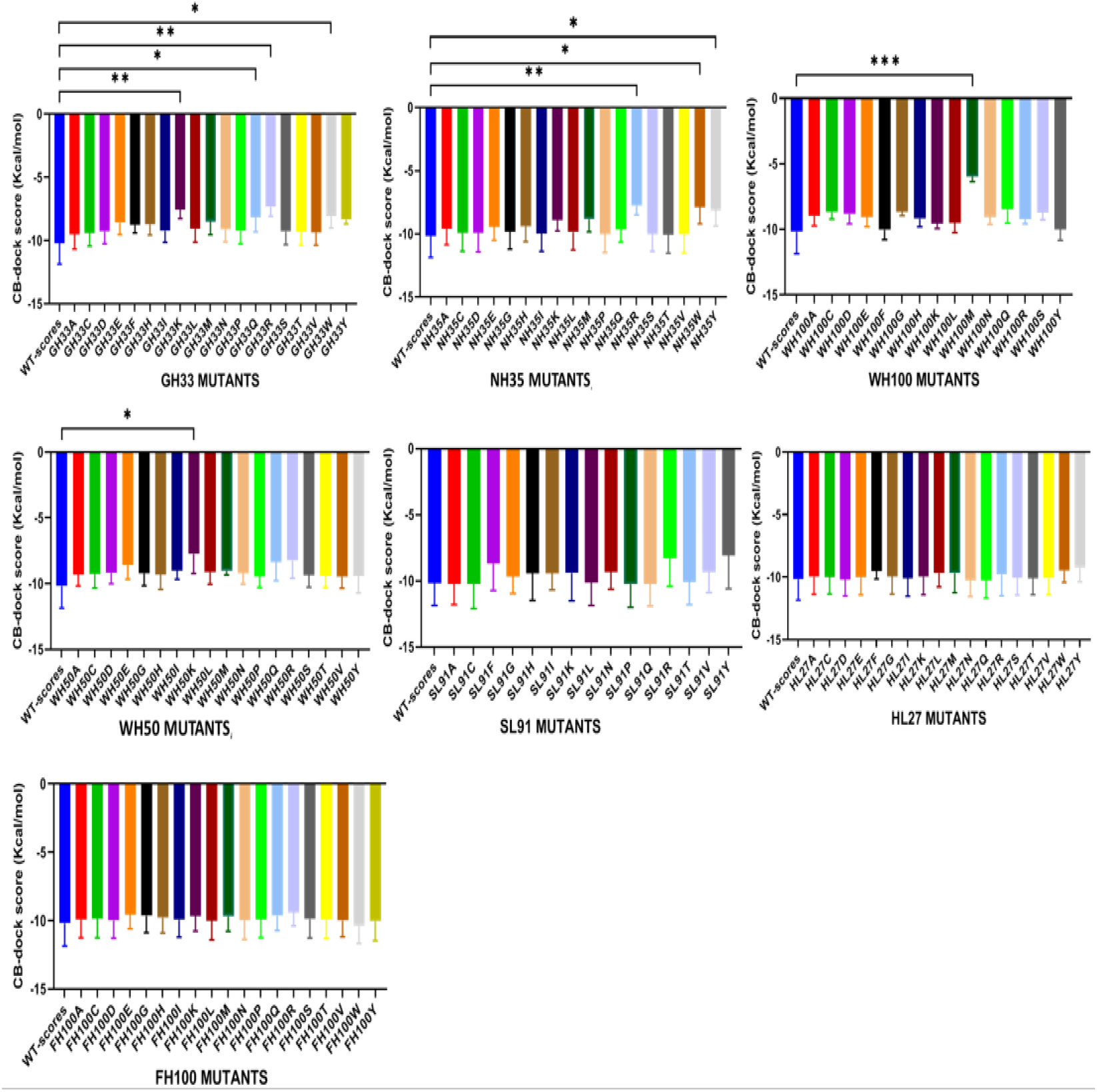
Docking scores. Selected residues were mutated to other amino acids, and their affinity was evaluated by docking in six structures and an average score was calculated. Statistically, significant differences are shown by * symbol.

**FIGURE 6.**
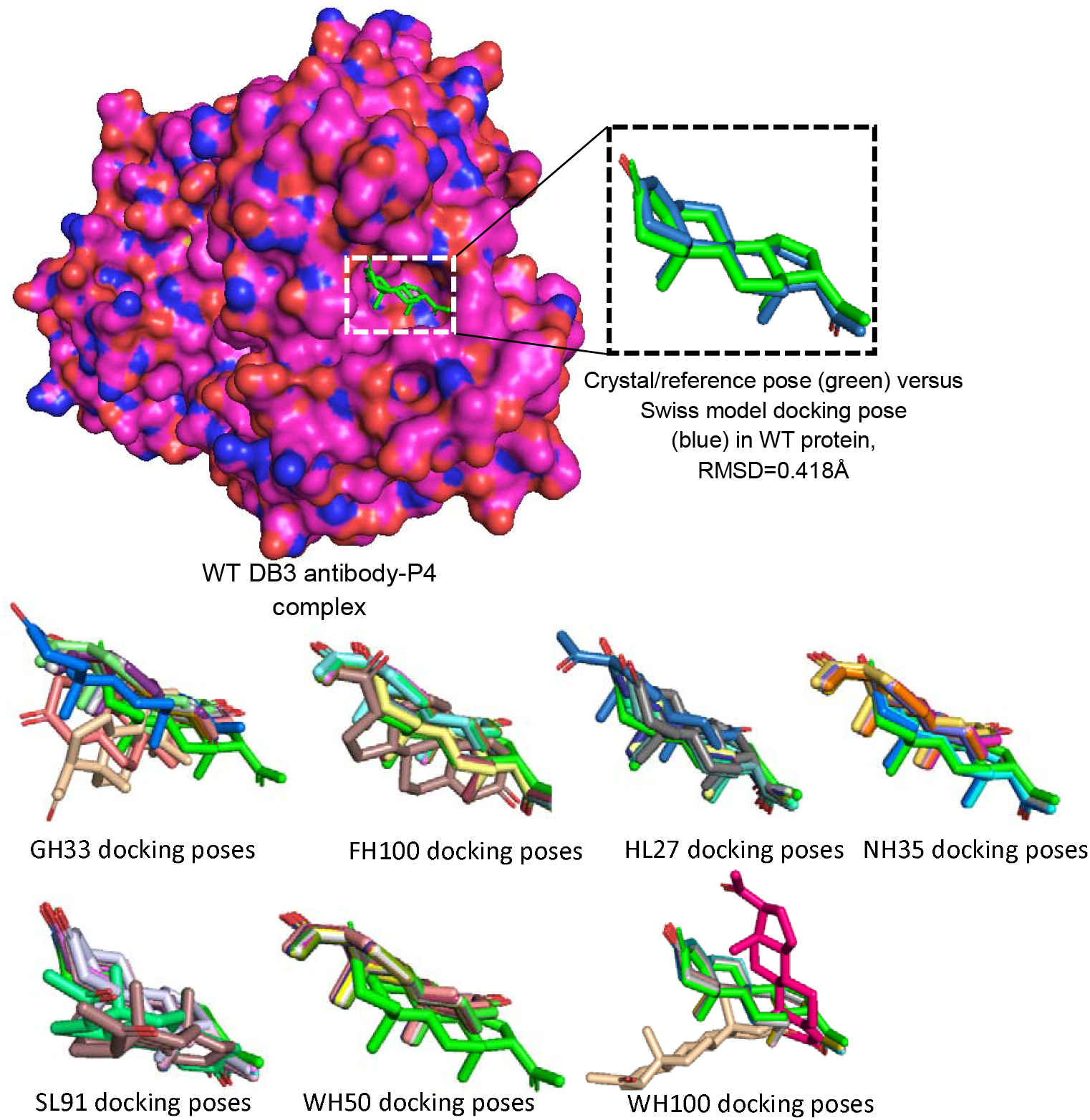
Study of binding site mutations on docking poses. Effect of mutations on binding pose was evaluated using the docking using 1dbb structure as a reference. GH33 docking poses were dissimilar, which agrees with the results of binding scores; however, the poses in the mutants of the other residues had some similarity with the WT pose. HL27 and SL91 mutants produced more WT-like docking poses and docking poses of WH50 mutants all were clustered in a 180 degree different conformer position where D-ring and A-ring locations were reversed entirely in the pocket. This might indicate that any mutation in the WH50 position can result in loss of specificity, as observed in NH35 mutants.

GH33K, GH33Q, GH33R, GH33W, NH35R, NH35W, NH35Y, WH100M, and WH50K mutations resulted in statistically more positive docking scores. WH100M was found to be the most impactful mutation, with a docking score of ∼5.5 kcal/mol. The presence of a higher number of statistically significant mutations of GH33 residue is in accordance with mutateX results, where mutations at this point were found to be energetically unfavourable (figure 4). Furthermore, the alanine scanning results agree with the docking results. Also, for the residues responsible for selective recognition of P4, i.e. NH35 and FH100, no statistically significant mutants of FH100 residue were found. However, some mutations in the NH35 position suggest that this residue might have a role in determining the binding affinity and the selectivity of DB3 anti-P4 antibody. In brief, no mutation was found to improve affinity in this step; however, as for NH35 position, some mutations could result in an inverse binding pose, indicating their effect on selectivity. As suggested by NH35 mutations, an inverse binding pose can result in binding other steroids to the same antibody. As mentioned earlier, the binding of the D-ring to the pocket ensures high selectivity for P4, and in the inverse binding pose, the D-ring of P4 molecule is not bound to the pocket, while other part of the molecule, which is highly similar to other steroid molecules could bind. The results of docking also indicate that mutations in the Heavy (H) chain are more likely to alter the affinity and or selectivity of the DB3 P4-antibody.

In some applications, selective recognition/biosensing might not be the case. For example, total cholesterol-derived steroids, i.e. mixed/crude steroid molecules, are monitored in samples of different origins such as environment^46^ and steroid bioprocesses^47^. In the steroid biotechnology industry, the discovery of new steroid-producing/metabolising microbes can also be facilitated via biomonitoring total steroid levels in the microbial culture media as a screening measure to speed up the strain discovery process^47^. In these cases, devising biosensing methods that are not selective among the steroid molecules but merely detect a group of steroids non-selectively can serve the purpose of steroid molecule screening. For reaching these mentioned aims, the first step is to design antibody mutants that have high affinity but that have lost their selectivity towards multiple cholesterol-derived steroid molecules eliminating the need to use multiple antibodies to measure each cholesterol-derived steroid individually.

In summary, this study section demonstrated that alanine scanning could identify important residues for each P4-binding protein; further modification of these residues can result in useful engineered versions of these proteins. Furthermore, alanine scanning results were cross validated using a combination of modelling and docking protocols, resulting in suggestions for designing putative anti-steroid anti-bodies based on the DB3 anti-P4 antibody.

After summarizing *in silico* studies on P4-bound pockets, the following points were found:

1. Selecting hydrophobic and non-polar amino acids in the pocket is crucial in designing or optimising P4-binding proteins. This fact agrees with the degree of hydrophobicity of the P4 molecule, with *logP10* (Partition Coefficient) Value of ∼4^48^, indicating that the binding pocket should contain enough hydrophobic amino acids to favour partitioning and interaction of P4 molecule with the protein. To support this fact, *in silico* mutations were performed on anti-DB3 antibody, and as evident from figure 5, most of the mutations that resulted in lower binding affinities are from amino acids with charged side chains.
2. Shape complementarity is another factor affecting the affinity and selectivity of pockets in steroid molecules^49^. The binding pocket should complement P4’s molecular shape to increase the affinity and selectivity of recognition. Nevertheless, recent studies suggest that the flexibility of the binding pocket can affect its complementarity, which should be taken into account by using molecular dynamics-based analysis of pocket shape^50^. In the current study, all structures of the same sequence were considered to approximate the docking results; however, future studies might require using molecular dynamics when doing the docking, which has not been done due to time constrains.
3. The findings showed that high-affinity anti-P4 binders usually have pocket sizes of around 500 Å3 (table 3). Previous studies have found that the internal volume of the pockets has a critical effect on the binding of ligands^51^. The internal volume of the pocket should be very close to the volume of P4, which is ∼ 447.86 Å^3^ and was calculated using the following formula: V=Mw/ρ, where V is volume, Mw is the molecular weight of P4, and ρ is the density of P4. The reason for this is to increase the selectivity of P4 over other hydrophobic molecules. As reported for the ancestral 3-keto steroid, a larger pocket results in non-specific interactions between the ligand and pocket and a diminished specificity for P4^52^. Therefore, the internal volume of the pocket should be very close to P4 volume to increase the binding specificity. Another factor to consider is that the volume of binding pockets can change due to the vibrational dynamics of proteins. Therefore, measuring pocket size at different time points using molecular dynamic simulation studies is recommended^50^ in future studies.
4. Some amino acids in the pocket can simultaneously form hydrophobic and/or hydrogen bonds with different groups, carbons and rings of P4. These interactions are required to increase the affinity and selectivity of P4 recognition. For instance, alanine and valine can solely form alkyl bonds; however, tryptophan and phenylalanine can form hydrogen, pi, sigma and alkyl bonds. As P4 is a steroid molecule and shares a considerable similarity with other steroids, it is necessary to design pockets that are highly P4-selective. Shape complementarity to the D ring and interaction of pocket residues with the D ring via hydrophobic and hydrogen bonds (C3=O group)^22^ are critical factors that should be considered to increase selectivity. In this study, it was found that NH35 residue of DB3 anti-P4 is important in both selectivity and affinity towards P4.
5. Increasing the overall hydrophobic interactions and hydrogen bonds between P4 and residues might improve the affinity of pockets by decreasing the binding energy.

As the data source of this study is the available resolved protein-P4 structures, the above main findings are based upon and reflect empirical experimental results. Therefore, the study of available experimental structures is recommended to improve the accuracy of any de novo computational approaches for designing small molecule binders. Moreover, these rules can be applied in library-based protein designing experiments to help decrease the initial size of in vitro mutants’ libraries, which can speed up the binder selection processes.

In future studies, it is possible to improve the accuracy of the main findings by including more parameters, such as molecular dynamics studies of P4-binding pockets. Furthermore, the field application of a newly designed protein requires considering binding kinetics, such as on-rates and off-rates^53^, which play essential roles in designing binders for use in competitive assays. These assays rely on the difference between the off-rates of competing molecules, i.e., manually added and sample P4 molecules.

### 3.3. In search of novel P4-binding protein scaffolds with favourable physicochemical properties using reverse docking

The final selected candidates from the target fishing tools are summarised in table 6. The candidates with Mw >25KDa were excluded from the final list of potential scaffolds. Receptors and enzymes were also not included among the final list of candidates. Lipocalin scaffolds were found to be the most abundant candidates suggested by the employed target fishing tools, followed by allosteric transcriptional factors (aTFs). The fibronectin family and laminin G-like domain were the third type of proteins, as the reverse docking tools suggested. Finding lipocalin as the principal scaffold in this study is in agreement with the previous reports on the potential of lipocalins for developing novel small molecule-binders due to the pocket geometries that enable them to bind to small molecules with high affinity and selectivity^19^. Lipocalin folds usually exhibit stable conformations with small sizes and monomeric structures^54^. The second group of potential candidates were aTFs whose activity is controlled by binding to small molecule ligands. aTFs have also been successfully engineered for the detection of small molecules^55^, including P4^56,57^ as reported in the literature, supporting the present in silico findings.

**Table 6.**
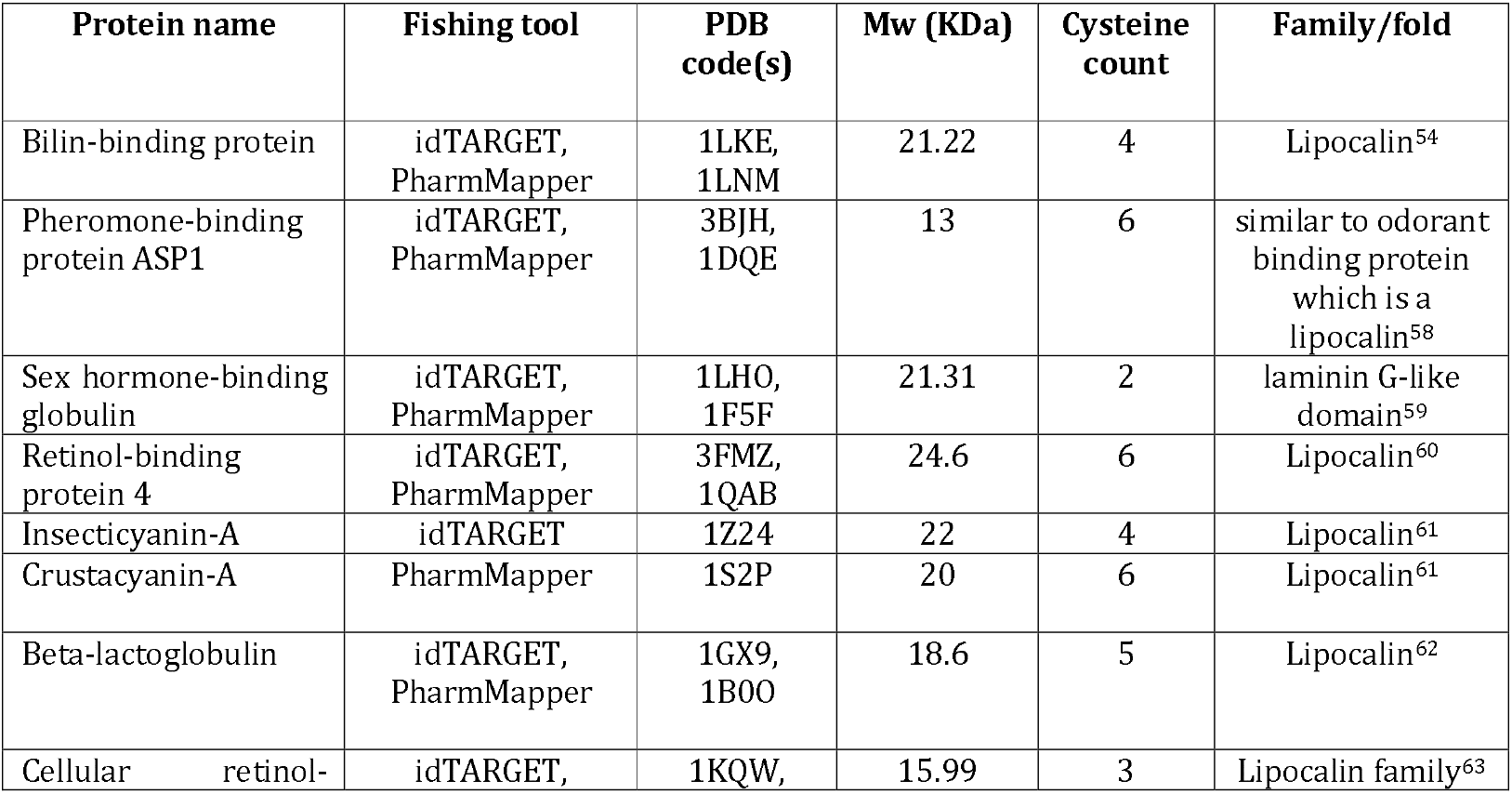
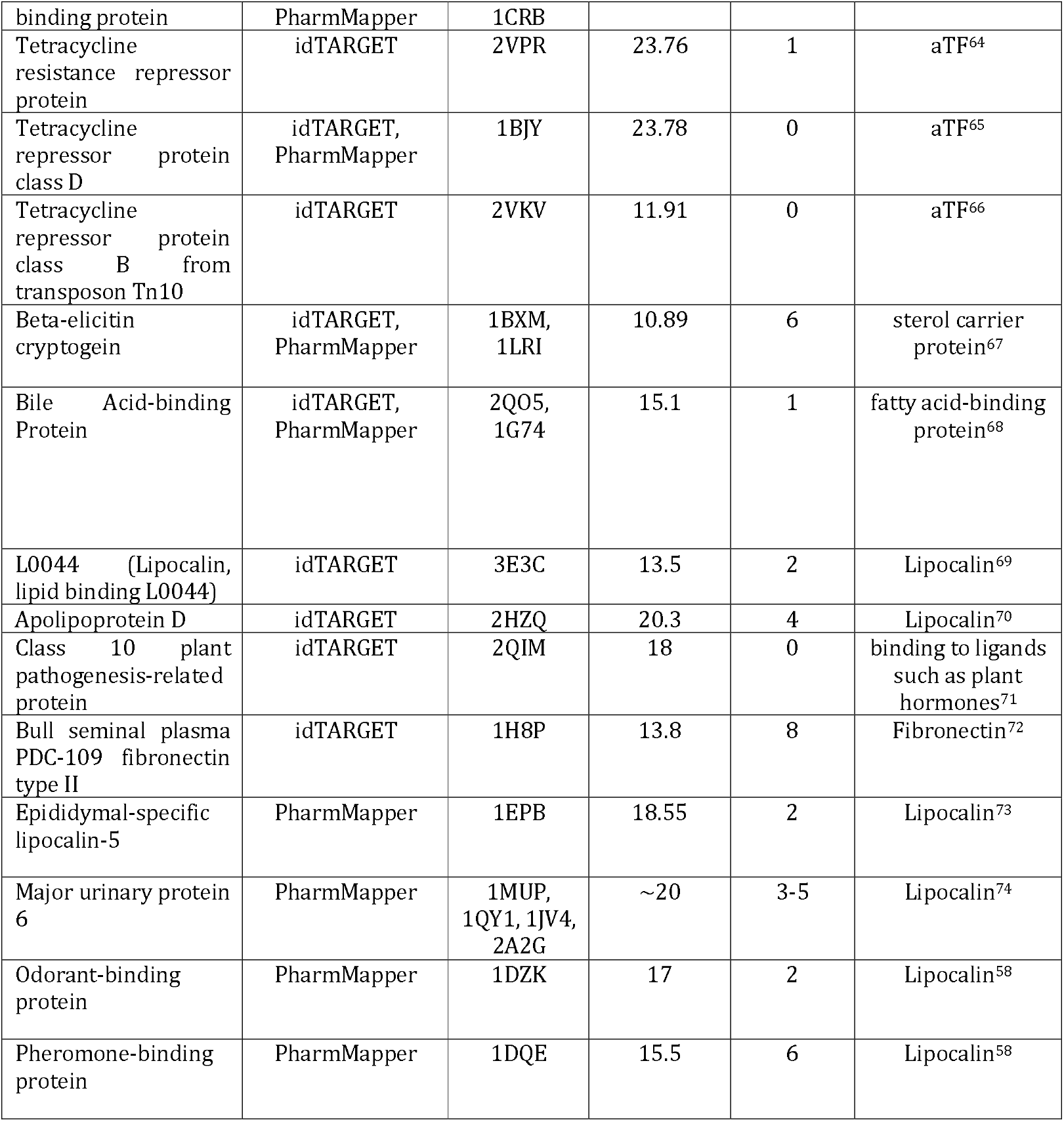
The final candidates suggested by performing reverse docking.

In brief, this study provided an overview of scaffolds for designing new binders and showed the possibility of fishing for novel scaffolds that can be engineered for P4 detection.

#### Correlation with Literature Data

For the anti-progesterone antibody DB3 (PDB: 1DBB), the predicted binding free energy (ΔG ≈ –9.8⍰kcal/mol) is moderately less favourable than the experimental estimate (ΔG ≈ –12.3⍰kcal/mol), calculated from the reported association constant (Ka ≈ 10^9^⍰M^−1^) using the thermodynamic relation ΔG = –RT⍰ln(Ka) at 298⍰K. This deviation is within the expected range for docking-based methods, which are optimised for relative ranking rather than precise thermodynamic accuracy. Computational alanine scanning identified TrpH100 as a critical contact residue, consistent with crystallographic data^75,76^.

A microbial allosteric transcription factor (aTF)-based biosensor has demonstrated sensitive detection of progesterone in artificial urine^56^. This functional outcome aligns with predicted binding energies in the range of –8 to –9⍰kcal/mol, supporting the suitability of aTFs as hormone-sensing scaffolds and validating the reverse docking approach employed.

Similarly, docking analyses of lipocalin scaffolds yielded binding energies between –7.5 and –8.8⍰kcal/mol, with pocket geometries that closely match known steroid-binding lipocalin structures. These results are consistent with prior engineering efforts that achieved nanomolar affinities for small hydrophobic ligands, including steroids^54^.

Taken together, the predicted binding energies, identified contact residues (e.g., TrpH100), and scaffold geometries exhibit strong agreement with experimental observations. This convergence supports the reliability of the computational workflow and the rationale behind scaffold and mutation selection for de novo progesterone binder design.

## 4. Conclusion

This study demonstrates the utility of *in silico* approaches, supported by experimentally reported data from the literature, for defining progesterone (P4) binding pocket features and identifying protein scaffolds capable of supporting these binding characteristics. Analysis of P4-binding pockets elucidated key geometric features, residue compositions, and interaction properties associated with high affinity and selectivity for P4. In addition, non-selective variants of an anti-P4 antibody were identified through *in silico* mutagenesis and molecular docking, revealing binding profiles compatible with multiple cholesterol-derived steroids. Furthermore, a reverse-docking strategy enabled the identification of alternative protein scaffolds exhibiting favourable physicochemical properties and predicted P4-binding capability.

Future work should focus on transferring the binding-pocket features identified in this study to rationally refine the pockets of selected scaffold candidates, followed by experimental validation of binding performance through *in vitro* and functional assays. Collectively, the outcomes of this study, together with subsequent validation efforts, represent an important step toward the development of next-generation progesterone and related small-molecule biosensing platforms based on custom-designed binders. Any future translation or commercialisation of such platforms will require the development of in-house binding reagents to support intellectual property protection and to minimise costs associated with large-scale production.

