## Supplementary material for "Designing High-Affinity Progesterone Binders: Pocket Analysis and Scaffold Selection": tables S1-S19

**¥Current Affiliations:**

a. Graduate School of Biomedical Engineering, Faculty of Engineering, University of New South Wales, Sydney 2052, Australia.
b. BioPoint Pty Ltd, Belrose 2085, Australia.

***Correspondence**:

**Table S1.** Biochemical properties of steroid receptors pockets.

| Name of protein  (PDB code) | Direction of ring interactions | Distribution of hydrogen bonds | Interpolated atomic charge | Hydrophobicity | Ionisability | Solvent accessible surface |
| --- | --- | --- | --- | --- | --- | --- |
|  | 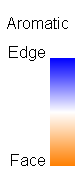 | 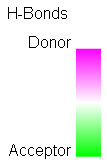 | 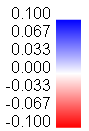 | 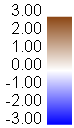 | 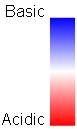 | 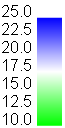 |
| Progesterone receptor ligand-binding domain (1a28) | 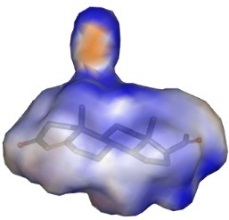 | 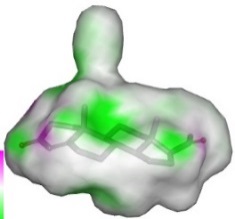 | 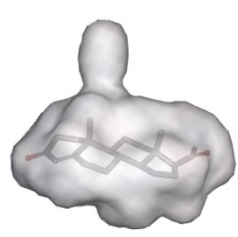 | 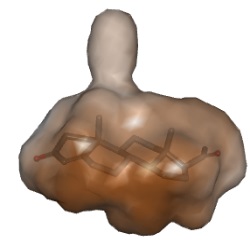 | 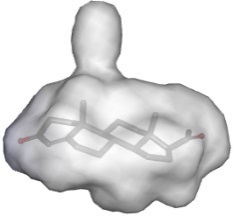 | 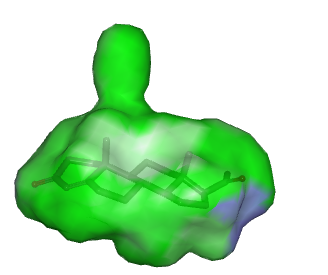 |
| Human mineralocorticoid receptor (1ya3) | 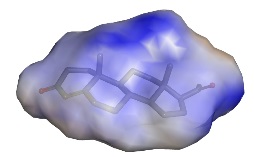 | 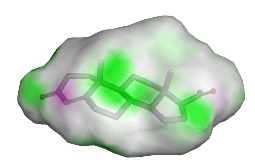 | 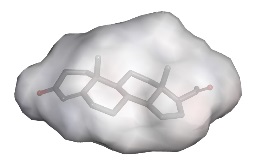 | 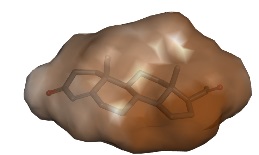 | 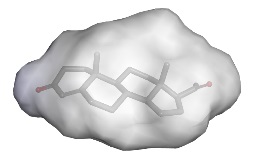 | 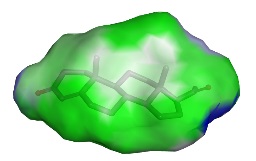 |
| Human mineralocorticoid receptor (2aa5) | 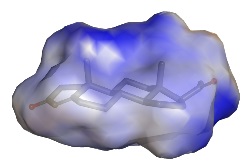 | 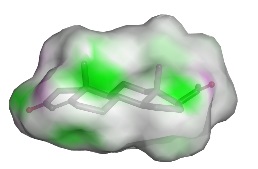 | 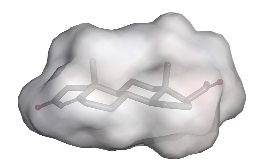 | 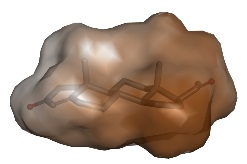 | 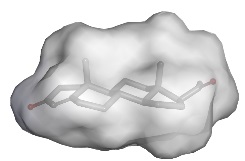 | 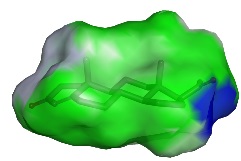 |
| Human mineralocorticoid receptor (2aa6) | 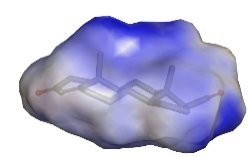 | 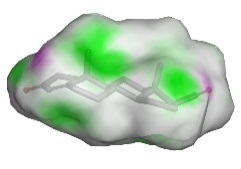 | 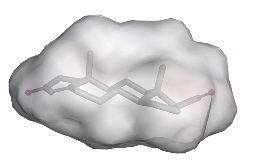 | 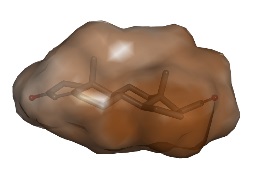 | 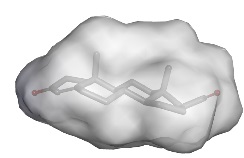 | 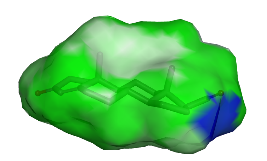 |
| Ancestral 3-keto steroid receptor (4fn9) | 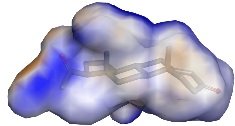 | 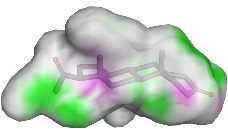 | 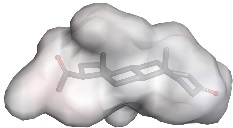 | 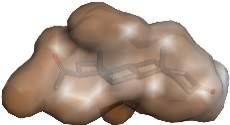 | 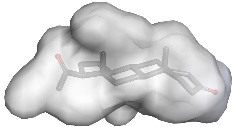 | 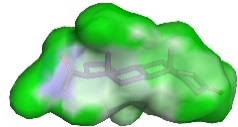 |
| Ancestral Ketosteroid Receptor (4ltw) | 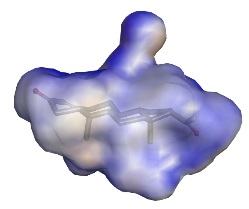 | 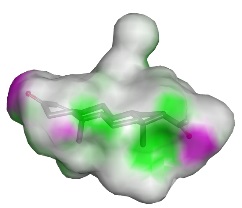 | 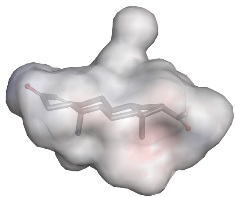 | 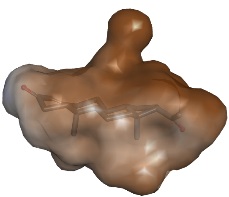 | 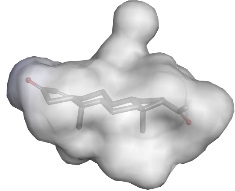 | 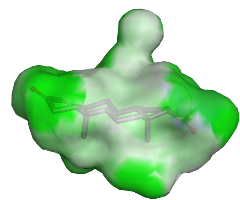 |

**Table S2.** Biochemical properties of anti-progesterone antibodies pockets.

| Name of protein  (PDB code) | Direction of ring interactions | Distribution of hydrogen bonds | Interpolated atomic charge | Hydrophobicity | Ionisability | Solvent accessible surface |
| --- | --- | --- | --- | --- | --- | --- |
|  | 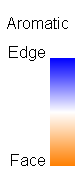 | 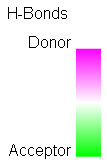 | 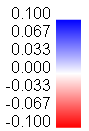 | 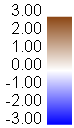 | 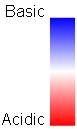 | 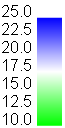 |
| DB3 anti-progesterone antibody (1dbb) | 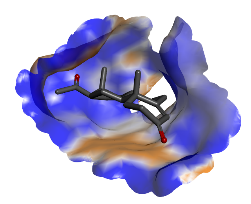 | 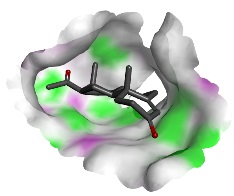 | 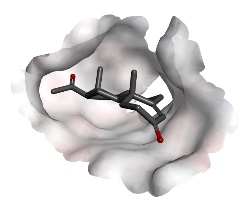 | 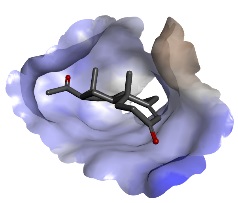 | 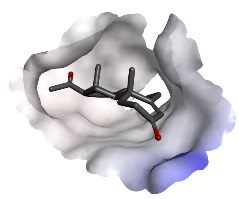 | 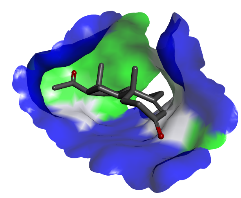 |
| DB3 anti-progesterone antibody (1dbm) | 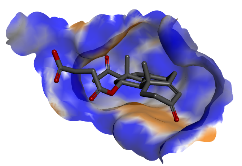 | 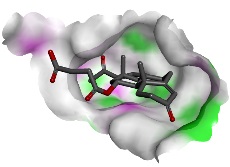 | 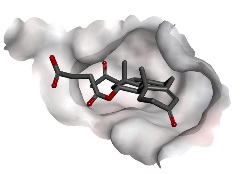 | 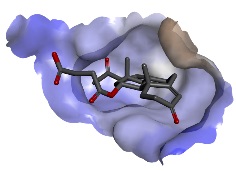 | 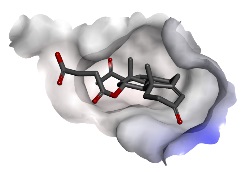 | 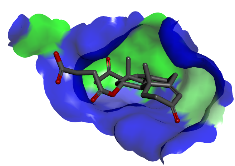 |
| 1E9 anti-progesterone antibody (2o5y) | 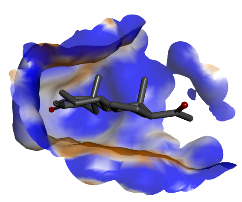 | 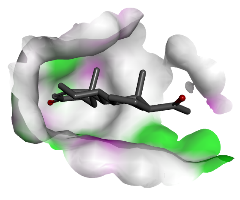 | 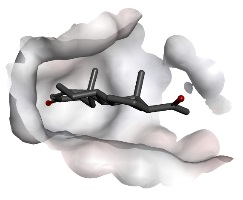 | 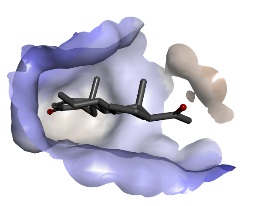 | 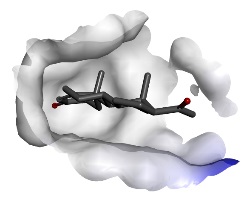 | 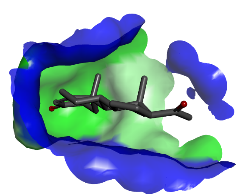 |

**Table S3.** Biochemical properties of steroid hydroxylase pockets.

| Name of protein  (PDB code) | Direction of ring interactions | Distribution of hydrogen bonds | Interpolated atomic charge | Hydrophobicity | Ionisability | Solvent accessible surface |
| --- | --- | --- | --- | --- | --- | --- |
|  | 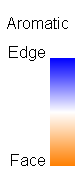 | 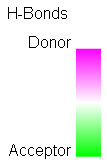 | 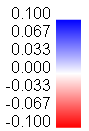 | 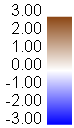 | 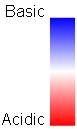 | 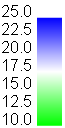 |
| Human steroidogenic cytochrome P450 (4nkx) | 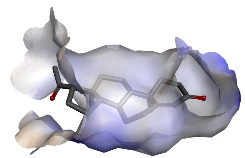 | 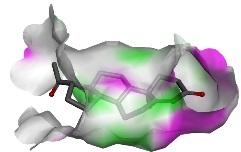 | 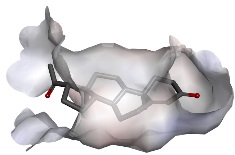 | 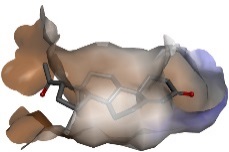 | 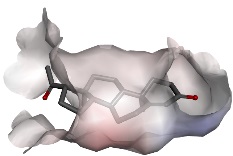 | 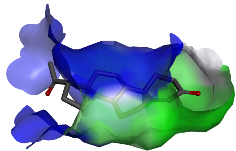 |
| Human steroidogenic cytochrome P450 (4nky) | 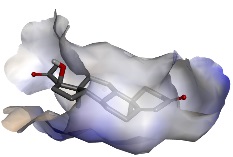 | 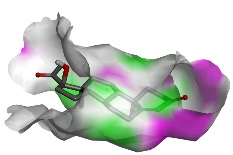 | 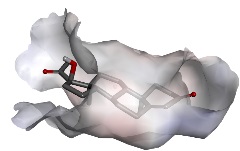 | 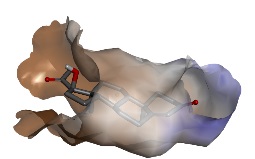 | 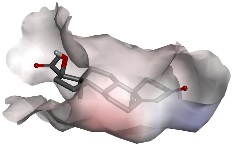 | 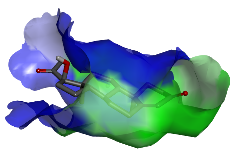 |
| Human steroidogenic cytochrome P450 (4nkz) | 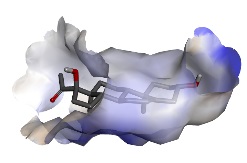 | 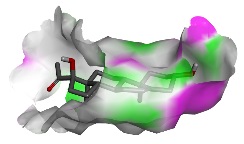 | 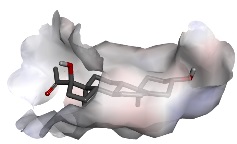 | 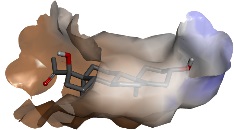 | 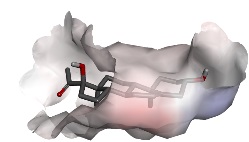 | 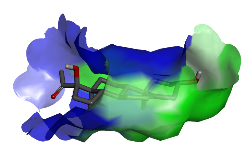 |

**Table S4.** Biochemical properties of reductase enzymes pockets.

| Name of protein  (PDB code) | Direction of ring interactions | Distribution of hydrogen bonds | Interpolated atomic charge | Hydrophobicity | Ionisability | Solvent accessible surface |
| --- | --- | --- | --- | --- | --- | --- |
|  | 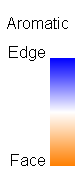 | 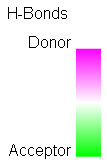 | 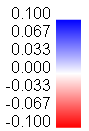 | 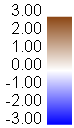 | 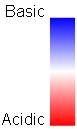 | 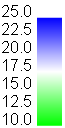 |
| Pentaerythritol tetranitrate reductase (1h60) | 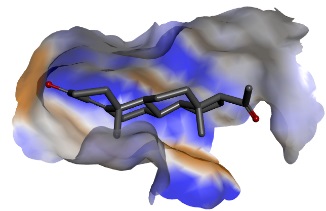 | 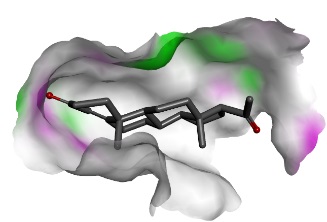 | 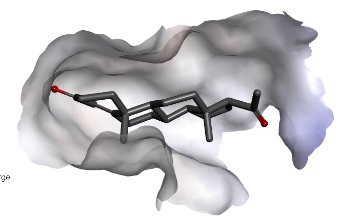 | 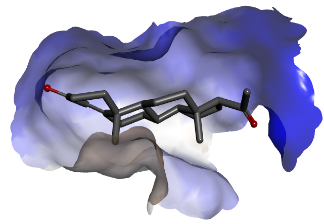 |  |  |
| Pentaerythritol tetranitrate reductase (2aba) |  |  |  |  |  |  |
| Human liver delta (4)-3-ketosteroid 5beta-reductase (3cot) |  |  |  |  |  |  |

**Table S5.** Biochemical properties of hydroxysteroid dehydrogenase enzymes pockets.

| Name of protein  (PDB code) | Direction of ring interactions | Distribution of hydrogen bonds | Interpolated atomic charge | Hydrophobicity | Ionisability | Solvent accessible surface |
| --- | --- | --- | --- | --- | --- | --- |
| Human 3-α-Hydroxysteroid Dehydrogenase Type 3 (4l1w) |  |  |  |  |  |  |
| Human 3-α-Hydroxysteroid Dehydrogenase Type 3 (4l1x) |  |  |  |  |  |  |
| Human 20-α-hydroxysteroid dehydrogenase (1mrq) |  |  |  |  |  |  |

**Table S6.** Biochemical properties of oxidoreductase enzymes pockets.

| Name of protein  (PDB code) | Direction of ring interactions | Distribution of hydrogen bonds | Interpolated atomic charge | Hydrophobicity | Ionisability | Solvent accessible surface |
| --- | --- | --- | --- | --- | --- | --- |
| Plantago Major multifunctional oxidoreductase (5mlm) |  |  |  |  |  |  |
| Plantago Major multifunctional oxidoreductase (6gsd) |  |  |  |  |  |  |

**Table S7.** Biochemical properties of steroid binding proteins pockets.

| Name of protein  (PDB code) | Direction of ring interactions | Distribution of hydrogen bonds | Interpolated atomic charge | Hydrophobicity | Ionisability | Solvent accessible surface |
| --- | --- | --- | --- | --- | --- | --- |
| Corticosteroid-binding globulin (4bb2) |  |  |  |  |  |  |
| α-1-antichymotrypsin variant NewBG-III (6hgk) |  |  |  |  |  |  |

**Table S8.** Biochemical properties of cytochrome P450 pockets.

| Name of protein  (PDB code) | Direction of ring interactions | Distribution of hydrogen bonds | Interpolated atomic charge | Hydrophobicity | Ionisability | Solvent accessible surface |
| --- | --- | --- | --- | --- | --- | --- |
| Cytochrome P450 CYP260A1(6f8c) |  |  |  |  |  |  |
| Cytochrome P450 CYP260A1 (6f88) |  |  |  |  |  |  |
| Cytochrome P450 3A4 (Cyp3A4 Monooxygenase) (5a1r) |  |  |  |  |  |  |
| Cytochrome P450 3A4 (Cyp3A4 Monooxygenase) (5a1p) |  |  |  |  |  |  |
| Human cytochrome P450 3A4 (1w0f) |  |  |  |  |  |  |
| Human Cytochrome P450 21A2 (4y8w) |  |  |  |  |  |  |
| Zebra fish cytochrome P450 17A2 (4r21) |  |  |  |  |  |  |
| Bacterial P450 monooxygenase (CYP154C5) (4j6c) |  |  |  |  |  |  |

**Table S9.** Biochemical properties of other progesterone-binding proteins pockets.

| Name of protein  (PDB code) | Direction of ring interactions | Distribution of hydrogen bonds | Interpolated atomic charge | Hydrophobicity | Ionisability | Solvent accessible surface |
| --- | --- | --- | --- | --- | --- | --- |
| Human apolipoprotein D (2hzq) |  |  |  |  |  |  |
| Engineered avidin (5lur) |  |  |  |  |  |  |

**Table S10.** (A): Details of biochemical interactions between progesterone and progesterone-binding steroid receptors.

| **Protein (PDB ID)** | **Involved amino acid residue (number)** | **Involved P4 site** | **Interaction type** | **Interaction distance (Å)** |
| --- | --- | --- | --- | --- |
| **Progesterone receptor ligand binding domain (1a28)** | ARG (766) | C_3_=O | conventional hydrogen bond | 2.77 |
|  | HOH (1004) | C_3_=O | water hydrogen bond | 3.09 |
|  | MET (759) | C19 | alkyl | 3.93 |
|  | TRP (755) | C19 | pi-alkyl | 5.44 |
|  | MET (756) | C19 | alkyl | 5.31 |
|  | MET (756) | C18 | alkyl | 5.45 |
|  | LEU (718) | C ring | alkyl | 5.06 |
|  | CYS (891) | C18 | alkyl | 3.76 |
|  | LEU (797) | D ring | alkyl | 4.96 |
|  | TYR (890) | D ring | pi-alkyl | 5.3 |
|  | CYS (891) | C_20_=O | carbon hydrogen bond | 3.3 |
| **Human mineralocorticoid receptor ligand-binding domain (1ya3)** | GLN 776 | C_3_=O | conventional hydrogen bond | 3.34 |
|  | LEU 810 | C19 | Alkyl | 4.26 |
|  | ALA 773 | C19 | Alkyl | 3.72 |
|  | MET 807 | C19 | Alkyl | 5.3 |
|  | MET 807 | B ring | Alkyl | 5.46 |
|  | MET 807 | C18 | Alkyl | 5.36 |
|  | CYS 942 | C18 | Alkyl | 3.68 |
|  | CYS 942 | C_20_=O | carbon hydrogen bond | 3.44 |
|  | LEU 769 | C ring | Alkyl | 5.36 |
|  | LEU 848 | D ring | Alkyl | 5.25 |
|  | LEU 938 | D ring | Alkyl | 4.97 |
|  | MET 845 | D ring | Alkyl | 4.27 |
| **Mineralocorticoid Receptor (2aa5)** | ARG 817 | C_3_=O | conventional hydrogen bond | 2.88 |
|  | TRP 806 | C19 | Pi-Alkyl | 5.14 |
|  | ALA 773 | C19 | Alkyl | 3.64 |
|  | MET 807 | B ring | Alkyl | 5.41 |
|  | MET 807 | C18 | Alkyl | 5.48 |
|  | LEU 769 | C ring | Alkyl | 5.08 |
|  | CYS 942 | C18 | Alkyl | 3.74 |
|  | CYS 942 | C_20_=O | carbon hydrogen bond | 3.23 |
|  | THR 945 | C_20_=O | conventional hydrogen bond | 2.72 |
|  | LEU 938 | D ring | Alkyl | 5.17 |
|  | MET 845 | D ring | Alkyl | 4.6 |
|  | PHE 941 | D ring | Pi-Alkyl | 5.27 |
|  | HOH 9 | C_3_=O | water conventional hydrogen bond | 3.02 |
|  | LEU 810 | C19 | Alkyl | 4.32 |
| **Mineralocorticoid Receptor (2aa6)** | ARG 817 | C_3_=O | conventional hydrogen bond | 2.8 |
|  | TRP 806 | C19 | Pi-Alkyl | 5.37 |
|  | ALA 773 | C19 | Alkyl | 3.61 |
|  | MET 807 | B ring | Alkyl | 5.43 |
|  | MET 807 | C18 | Alkyl | 5.44 |
|  | MET 807 | C19 | Alkyl | 5.32 |
|  | LEU 769 | C ring | Alkyl | 5.04 |
|  | CYS 942 | C18 | Alkyl | 3.82 |
|  | CYS 942 | C_20_=O | carbon hydrogen bond | 3.17 |
|  | THR 945 | C_20_=O | conventional hydrogen bond | 2.98 |
|  | LEU 938 | D ring | Alkyl | 5.14 |
|  | MET 845 | D ring | Alkyl | 4.34 |
|  | PHE 941 | D ring | Pi-Alkyl | 5.27 |
|  | HOH 9 | C_3_=O | water conventional hydrogen bond | 3.18 |
|  | LEU 810 | C19 | Alkyl | 4.32 |
| **Ancestral 3-keto steroid receptor (4fn9)** | ARG 82 | C_3_=O | conventional hydrogen bond | 2.84 |
|  | TRP 71 | C19 | pi-alkyl | 5.4 |
|  | MET 75 | C19 | alkyl | 4.12 |
|  | MET 72 | B ring | alkyl | 5.3 |
|  | MET 72 | C19 | alkyl | 4.7 |
|  | ALA 38 | C19 | alkyl | 4.19 |
|  | ALA 38 | C ring | alkyl | 5.24 |
|  | THR 210 | C_20_=O | conventional hydrogen bond | 2.98 |
|  | MET 110 | D ring | alkyl | 4.66 |
|  | LEU 203 | D ring | alkyl | 5.21 |
|  | PHE 206 | D ring | pi-alkyl | 5.25 |
|  | LEU 34 | C ring | alkyl | 4.38 |
|  | LEU 34 | B ring | alkyl | 5.47 |
|  | ALA 76 | B ring | alkyl | 5.4 |
|  | GLN 41 | C_3_=O | conventional hydrogen bond | 2.99 |
|  | HOH 401 | C_3_=O | water conventional hydrogen bond | 3.28 |
| **Ancestral Ketosteroid Receptor (4ltw)** | CYS 207 | C_20_=O | carbon hydrogen bond | 2.31 |
|  | CYS 207 | D ring | alkyl | 5.09 |
|  | MET 110 | D ring | alkyl | 4.85 |
|  | PHE 206 | D ring | pi-alkyl | 5.45 |
|  | LEU 203 | D ring | alkyl | 5.02 |
|  | MET 72 | C19 | alkyl | 5.33 |
|  | ARG 82 | C_3_=O | conventional hydrogen bond | 2.46 |
|  | HOH 401 | C_3_=O | water conventional hydrogen bond | 3.17 |
|  | GLN 41 | C_3_=O | conventional hydrogen bond | 2.45 |
|  | MET 75 | C19 | alkyl | 4.16 |
|  | ALA 38 | C19 | alkyl | 3.91 |
|  | ALA 38 | C ring | alkyl | 5.45 |
|  | LEU 34 | C ring | alkyl | 4.81 |

**Table S10.** (B): Consensus binding residues of the pocket and interactions between P4 and the steroid receptors.

| Involved amino acid residue | Parent protein(s) | Average Residue number per pocket | Involved P4 site(s) | Interaction type | Average Interaction distance (Å) |
| --- | --- | --- | --- | --- | --- |
| ARG | All receptors excluding Human mineralocorticoid receptor ligand-binding domain (1ya3) | 1 | C_3_=O | conventional hydrogen bond | 2.75 |
| MET | All receptors | 3.5 | C19, C18, D ring, B ring | alkyl | 4.28 |
| TRP | All except Ancestral Ketosteroid Receptor (4ltw) and Human mineralocorticoid receptor ligand-binding domain (1ya3) | 0.67 | C19 | pi-alkyl | 5.34 |
| LEU | All receptors | 2.6 | C ring, D ring, B ring, C19 | alkyl | 4.93 |
| CYS | All receptors excluding Ancestral 3-keto steroid receptor (4fn9) | 0.84 | C18, C_20_=O, D ring | carbon hydrogen bond, alkyl | 3.09 (for carbon hydrogen bond), 4.018 (for alkyl bond) |
| ALA | All receptors excluding Progesterone receptor ligand binding domain (1a28) | 1.34 | C19, C ring, B ring | alkyl | 4.4 |
| GLN | All receptors excluding Progesterone receptor ligand binding domain (1a28) and Mineralocorticoid Receptor (2aa5 and 2aa6) | 0.5 | C_3_=O | conventional hydrogen bond | 2.92 |
| PHE | All receptors excluding Progesterone receptor ligand binding domain (1a28) and Human mineralocorticoid receptor ligand-binding domain (1ya3) | 0.66 | D ring | pi-alkyl | 5.31 |

**Table S11**. (A): Details of biochemical interactions between progesterone and anti-progesterone antibodies.

| **Protein (PDB ID)** | | **Involved amino acid residue (number)** | **Involved P4 site** | **Interaction type** | **Interaction distance (Å)** |
| --- | --- | --- | --- | --- | --- |
| **Anti-progesterone DB3 antibody (1dbb)** | VAL, L chain (94) | | C19 | alkyl | 4.5 |
|  | TRP, H chain (50) | | C ring | pi-alkyl | 4.71 |
|  | TRP, H chain (50) | | C18 | pi-sigma | 3.91 |
|  | TRP, H chain (50) | | C18 | pi-sigma | 3.7 |
|  | ASN, H chain (35) | | C_20_=O | conventional hydrogen bond | 3.17 |
|  | PHE, H chain (100b) | | D ring | pi-alkyl | 5.24 |
| **Anti-progesterone DB3 antibody (1dbm)** | VAL, L chain (94) | | B ring | alkyl | 5.34 |
|  | PRO, L chain (96) | | B ring | alkyl | 5.36 |
|  | TRP, H chain (100) | | B ring | pi-alkyl | 4.64 |
|  | TRP, H chain (100) | | B ring | pi-alkyl | 4.59 |
|  | TRP, H chain (100) | | D ring | pi-alkyl | 4.61 |
|  | ASN, H chain (35) | | C_20_=O | conventional hydrogen bond | 3.4 |
|  | GLY, H chain (33) | | C_20_=O | carbon hydrogen bond | 3.25 |
|  | TRP, H chain (50) | | C_20_=O | conventional hydrogen bond | 2.81 |
|  | TRP, H chain (50) | | C18 | pi-sigma | 3.44 |
|  | TRP, H chain (50) | | C18 | pi-sigma | 3.61 |
|  | TYR, H chain (97) | | C21 | pi-sigma | 3.5 |
| **1E9 anti-progesterone antibody (2o5y)** | TRP, H chain 47 | | C19 | pi-alkyl | 5.18 |
|  | TRP, H chain 50 | | B ring | pi-sigma | 3.9 |
|  | TRP, H chain 50 | | B ring | pi-alkyl | 4.86 |
|  | TRP, H chain 50 | | C19 | pi-sigma | 3.62 |
|  | TRP, H chain 50 | | C18 | pi-alkyl | 5.5 |
|  | TRP, H chain 100 | | B ring | pi-alkyl | 4.95 |
|  | TRP, H chain 100 | | C ring | pi-sigma | 3.97 |
|  | TRP, H chain 100 | | D ring | pi-alkyl | 4.73 |
|  | TRP, H chain 100 | | D ring | pi-alkyl | 5.01 |

**Table S11.** (B): Consensus binding residues of the pocket and interactions between progesterone and the anti-P4 antibodies.

| Involved amino acid residue | Parent protein(s) | Average Residue number per pocket | Involved P4 site(s) | Interaction type | Average Interaction distance (Å) |
| --- | --- | --- | --- | --- | --- |
| VAL (Light chain) | anti-progesterone DB3 antibody (1dbb and 1dbm) | 0.67 | C19, B ring | alkyl | 4.92 |
| TRP (Heavy chain) | all anti-P4 antibodies | 2 | B ring, C ring, D ring, C18, C19, C_20_=O | pi-alkyl, pi-sigma, conventional hydrogen bond | 4.84 (pi-alkyl bond), 3.74 (pi-sigma), 2.81 (conventional hydrogen bond) |
| ASN (Heavy chain) | anti-progesterone DB3 antibody (1dbb and 1dbm) | 0.67 | C_20_=O | conventional hydrogen bond | 3.28 |

**Table S12.** (A): Details of biochemical interactions between progesterone and steroid hydroxylase enzymes

| **Protein (PDB ID)** | **Involved amino acid residue (number)** | | **Involved P4 site** | **Interaction type** | **Interaction distance (Å)** |
| --- | --- | --- | --- | --- | --- |
| **Human steroidogenic cytochrome P450 17A1 mutant A105L (4nkx)** | | ALA 113 | D ring | alkyl | 3.7 |
|  |  | PHE 114 | D ring | pi-alkyl | 5.4 |
|  |  | ALA 302 | D ring | alkyl | 3.86 |
|  |  | ALA 302 | C ring | alkyl | 4.36 |
|  |  | ASN 202 | C_3_=O | conventional hydrogen bond | 2.82 |
| **Human steroidogenic cytochrome P450 17A1 mutant A105L (4nky)** | | HOH 740 | C_3_=O | water conventional hydrogen bond | 3.35 |
|  |  | HOH 734 | C_3_=O | water conventional hydrogen bond | 3.34 |
|  |  | LEU 105 | C19 | alkyl | 5 |
|  |  | LEU 209 | C19 | alkyl | 5.19 |
|  |  | VAL 482 | C19 | alkyl | 4.61 |
|  |  | ALA 302 | D ring | alkyl | 3.95 |
|  |  | ALA 302 | C ring | alkyl | 4.52 |
|  |  | ALA 113 | C ring | alkyl | 3.8 |
| **Human steroidogenic cytochrome P450 17A1 mutant A105L (4nkz)** | | PHE 114 | D ring | pi-alkyl | 5.19 |
|  |  | ALA 113 | D ring | alkyl | 3.4 |
|  |  | ALA 302 | D ring | alkyl | 4.36 |
|  |  | ALA 302 | C ring | alkyl | 4.45 |
|  |  | ALA 302 | B ring | alkyl | 4.57 |
|  |  | LEU 105 | C19 | alkyl | 5.27 |
|  |  | LEU 209 | C19 | alkyl | 5.3 |
|  |  | ILE 205 | A ring | alkyl | 5.5 |
|  |  | ASN 202 | C_3_-OH | conventional hydrogen bond | 2.31 |
|  |  | VAL 482 | C19 | alkyl | 4.6 |

**Table S12.** (B): Consensus binding residues of the pocket and interactions between progesterone and the steroid hydroxylase enzymes.

| Involved amino acid residue | Parent protein(s) | Average Residue number per pocket | Involved P4 site(s) | Interaction type | Average Interaction distance (Å) |
| --- | --- | --- | --- | --- | --- |
| ALA | all | 2 | B, C and D rings | alkyl | 4.1 |
| PHE | human steroidogenic cytochrome P450 17A1 mutant A105L (4nkx and 4nkz) | 0.67 | D ring | Pi-alkyl | 5.3 |
| ASN | human steroidogenic cytochrome P450 17A1 mutant A105L (4nkx and 4nkz) | 0.67 | C_3_=O | conventional hydrogen bond | 2.57 |
| LEU | human steroidogenic cytochrome P450 17A1 mutant A105L (4nky and 4nkz) | 0.67 | C19 | alkyl | 5.19 |
| VAL | human steroidogenic cytochrome P450 17A1 mutant A105L (4nky and 4nkz) | 0.67 | C19 | alkyl | 4.6 |

**Table S13**. (A): Details of biochemical interactions between progesterone and reductase enzymes.

| **Protein (PDB ID)** | | **Involved amino acid residue (number)** | **Involved P4 site** | **Interaction type** | **Interaction distance (Å)** |
| --- | --- | --- | --- | --- | --- |
| **Pentaerythritol tetranitrate reductase (1h60)** | HIS (184) | | C_3_=O | conventional hydrogen bond | 2.76 |
|  | LEU (275) | | C19 | alkyl | 4.88 |
|  | FMN (401) | | C19 | pi-sigma | 3.43 |
|  | FMN (401) | | C19 | pi-sigma | 3.88 |
|  | FMN (401) | | B ring | pi-alkyl | 4.99 |
|  | FMN (401) | | B ring | Pi-Alkyl | 4.96 |
|  | TYR (68) | | B ring | Pi-Alkyl | 5.44 |
|  | TYR (68) | | D ring | Pi-Alkyl | 5.48 |
|  | HOH (2667) | | C_20_=O | water hydrogen bond (conventional) | 2.9 |
|  | HOH (2666) | | C_20_=O | water hydrogen bond (conventional) | 3.01 |
|  | TYR (351 | | D ring | Pi-Alkyl | 4.74 |
| **Reduced pentaerythritol tetranitrate reductase (2aba)** | HIS 184 | | C_3_=O | conventional hydrogen bond | 2.87 |
|  | LEU 275 | | C19 | Alkyl | 5.28 |
|  | TYR 68 | | B ring | Pi-Alkyl | 5.26 |
|  | TYR 68 | | D ring | Pi-Alkyl | 5.22 |
|  | ARG 142 | | C_20_=O | conventional hydrogen bond | 2.77 |
|  | ARG 142 | | C_20_=O | conventional hydrogen bond | 2.83 |
|  | HOH 1757 | | C_20_=O | water hydrogen bond | 2.99 |
|  | TYR 351 | | C18 | Pi-Alkyl | 4.27 |
|  | TYR 351 | | D ring | Pi-Alkyl | 4.8 |
| **Human liver-Δ-(4)-3-ketosteroid 5β-reductase (akr1d1) (3cot)** | TYR 58 | | 3C=O | conventional hydrogen bond | 2.61 |
|  | VAL 309 | | C19 | alkyl | 5.16 |
|  | LEU 311 | | C19 | alkyl | 4.87 |
|  | TRP 230 | | C19 | pi-alkyl | 4.9 |
|  | TRP 230 | | B ring | pi-alkyl | 4.69 |
|  | TRP 230 | | D ring | pi-alkyl | 4.37 |
|  | TRP 230 | | C18 | pi-alkyl | 5.17 |
|  | TRP 230 | | C18 | pi-alkyl | 4.44 |
|  | TYR 132 | | C ring | pi-alkyl | 5.31 |
|  | TYR 26 | | B ring | pi-alkyl | 4.93 |

**Table S13**. (B): Consensus binding residues of the pocket and interactions between progesterone and the reductase enzymes.

| Involved amino acid residue | Parent protein(s) | Average Residue number per pocket | Involved P4 site(s) | Interaction type | Average Interaction distance (Å) |
| --- | --- | --- | --- | --- | --- |
| HIS | Pentaerythritol tetranitrate reductase (1h60) and Reduced pentaerythritol tetranitrate reductase (2aba) | 0.67 | C3=O | conventional hydrogen bond | 5.63 |
| LEU | All | 1 | C19 | alkyl | 5.01 |
| TYR | All | 2.34 | B ring, C ring, D ring, C18,  C3=O | pi-alkyl, conventional hydrogen bond | 5.02 (for pi-alkyl bond), 2.61 (for conventional hydrogen bond) |

**Table S14**. (A): Details of biochemical interactions between progesterone and hydroxysteroid dehydrogenase enzymes.

| **Protein (PDB ID)** | | **Involved amino acid residue (number)** | **Involved P4 site** | **Interaction type** | **Interaction distance (Å)** |
| --- | --- | --- | --- | --- | --- |
| **Human 3-alpha Hydroxysteroid Dehydrogenase Type 3 (4l1w)** | TRP 227 | | C19 | pi-alkyl | 5.03 |
|  | TRP 227 | | C19 | pi-alkyl | 4.94 |
|  | TRP 227 | | B ring | pi-alkyl | 4.71 |
|  | TRP 227 | | B ring | pi-alkyl | 5.42 |
|  | TRP 227 | | D ring | pi-alkyl | 4.82 |
|  | TRP 227 | | C18 | pi-sigma | 3.76 |
|  | TYR 24 | | C ring | pi-alkyl | 5.13 |
|  | TYR 24 | | C18 | pi-sigma | 3.64 |
|  | VAL 54 | | C ring | alkyl | 5.35 |
|  | LEU 308 | | D ring | alkyl | 4.41 |
| **Human 3-α-hydroxysteroid dehydrogenase type 3 V54L Mutant (4l1x)** | HOH 749 | | C_3_=O | water conventional hydrogen bond | 2.8 |
|  | LEU 54 | | B ring | alkyl | 5.05 |
|  | LEU 54 | | C ring | alkyl | 4.77 |
|  | TRP 227 | | C19 | pi-alkyl | 5.15 |
|  | TRP 227 | | C19 | pi-alkyl | 4.87 |
|  | TRP 227 | | B ring | pi-alkyl | 4.6 |
|  | TRP 227 | | B ring | pi-alkyl | 4.95 |
|  | TRP 227 | | D ring | pi-alkyl | 4.98 |
|  | TRP 227 | | D ring | pi-alkyl | 5.3 |
|  | TRP 227 | | C18 | pi-sigma | 3.88 |
|  | TYR 24 | | C ring | pi-alkyl | 4.72 |
|  | LEU 308 | | D ring | alkyl | 4.52 |
| **Human 20-α-hydroxysteroid dehydrogenase (1mrq)** | TRP 227 | | B ring | Pi-Alkyl | 4.6 |
|  | TRP 227 | | B ring | Pi-sigma | 3.6 |
|  | TRP 227 | | C19 | Pi-Alkyl | 4.38 |
|  | TRP 227 | | C19 | Pi-Alkyl | 4.66 |
|  | TYR 24 | | C ring | Pi-Sigma | 3.98 |
|  | TYR 24 | | C21 | Pi-Sigma | 3.94 |
|  | TYR 55 | | C21 | Pi-Alkyl | 4.33 |
|  | HIS 222 | | C21 | Pi-Alkyl | 5.2 |
|  | HIS 222 | | C_20_=O | conventional hydrogen bond | 2.91 |
|  | LEU 308 | | D ring | Alkyl | 5.29 |
|  | LEU 306 | | D ring | Alkyl | 5.12 |
|  | LEU 54 | | C ring | Alkyl | 4.6 |
|  | LEU 54 | | B ring | Alkyl | 5.19 |

**Table S14.** (B): Consensus binding residues of the pocket and interactions between progesterone and the hydroxysteroid dehydrogenase enzymes.

| Involved amino acid residue | Parent protein(s) | Average Residue number per pocket | Involved P4 site(s) | Interaction type | Average Interaction distance (Å) |
| --- | --- | --- | --- | --- | --- |
| TRP | all | 1 | C18, C19, B ring, D ring | pi-alkyl, pi-sigma | 4.89 (for pi-alkyl bond), 3.75 (for pi-sigma) |
| TYR | all | 1.34 | C18, C21, C ring | pi-alkyl, pi-sigma | 4.58 (for pi-alkyl bond), 3.85 (for pi-sigma bond) |
| LEU | all | 2 | B ring, C ring, D ring | alkyl | 4.87 |

**Table S15**. (A): Details of biochemical interactions between progesterone and oxidoreductase enzymes.

| **Protein (PDB ID)** | **Involved amino acid residue (number)** | | **Involved P4 site** | **Interaction type** | **Interaction distance (Å)** |
| --- | --- | --- | --- | --- | --- |
| **Plantago Major multifunctional oxidoreductase V150M mutant (5mlm)** | | CYS 352 | C18 | alkyl | 4.34 |
|  |  | CYS 352 | D ring | alkyl | 3.08 |
|  |  | ARG 146 | B ring | alkyl | 5.29 |
|  |  | LYS 147 | B ring | alkyl | 5.25 |
|  |  | LYS 147 | C19 | alkyl | 3.95 |
|  |  | LYS 147 | C_3_=O | conventional hydrogen bond | 2.58 |
|  |  | MET 150 | B ring | alkyl | 4.82 |
|  |  | MET 150 | C19 | alkyl | 3.55 |
|  |  | ILE 156 | C19 | alkyl | 4 |
|  |  | VAL 347 | C ring | alkyl | 4.43 |
|  |  | VAL 347 | D ring | alkyl | 5.15 |
|  |  | ILE 350 | C ring | alkyl | 4.62 |
|  |  | ILE 350 | C18 | alkyl | 4.2 |
|  |  | PRO 353 | C18 | alkyl | 4.61 |
| Plantago Major multifunctional oxidoreductase (6gsd) | | PRO 353 | D ring | alkyl | 5.08 |
|  |  | CYS 352 | D ring | alkyl | 3.81 |
|  |  | ARG 146 | B ring | alkyl | 4.81 |
|  |  | VAL 150 | C19 | alkyl | 5.37 |
|  |  | LYS 147 | B ring | alkyl | 5.36 |
|  |  | LYS 147 | C19 | alkyl | 4.06 |
|  |  | ILE 156 | C19 | alkyl | 4.53 |
|  |  | VAL 347 | C ring | alkyl | 4.8 |
|  |  | ILE 350 | C ring | alkyl | 5.47 |
|  |  | ILE 350 | C18 | alkyl | 4.2 |

**Table S15**. (B): Consensus binding residues of the pocket and interactions between progesterone and the oxidoreductase enzymes.

| Involved amino acid residue | Parent protein(s) | Average Residue number per pocket | Involved P4 site(s) | Interaction type | Average Interaction distance (Å) |
| --- | --- | --- | --- | --- | --- |
| CYS | both | 1 | C18, D ring | alkyl | 3.74 |
| ARG | both | 1 | B ring | alkyl | 5.01 |
| LYS | both | 1 | C19, B ring, C3=O | alkyl, conventional hydrogen bond | 2.58 (for conventional hydrogen bond), 4.66 (for alkyl bond) |
| ILE | both | 2 | C18, C19, C ring | alkyl | 4.54 |
| VAL | both | 2 | C19, C ring, D ring | alkyl | 4.94 |
| PRO | both | 1 | C18, D ring | alkyl | 4.85 |

**Table S16.** (A): Details of biochemical interactions between progesterone and steroid binding proteins.

| **Protein (PDB ID)** | | **Involved amino acid residue (number)** | **Involved P4 site** | **Interaction type** | **Interaction distance (Å)** |
| --- | --- | --- | --- | --- | --- |
| **Corticosteroid-binding globulin (4bb2)** | HOH, B chain 2009 | | C_3_=O | water conventional hydrogen bond | 2.89 |
|  | ARG, A chain 260 | | C19 | alkyl | 4.16 |
|  | ILE, A chain 263 | | B ring | alkyl | 5.16 |
|  | ILE, A chain 263 | | C18 | alkyl | 5.42 |
|  | ILE, A chain 263 | | C19 | alkyl | 4.19 |
|  | TRP, B chain 371 | | A ring | pi-sigma | 3.88 |
|  | TRP, B chain 371 | | B ring | pi-alkyl | 4.88 |
|  | TRP, B chain 371 | | Between B and C rings (C9) | pi-sigma | 3.62 |
|  | TRP, B chain 371 | | C ring | pi-alkyl | 5.13 |
|  | TRP, B chain 371 | | C ring | pi-alkyl | 4.14 |
|  | TRP, B chain 371 | | D ring | pi-alkyl | 5.06 |
|  | HOH, A chain 2060 | | C_20_=O | water conventional hydrogen bond | 2.83 |
|  | GLN, A chian 232 | | C_20_=O | conventional hydrogen bond | 3.19 |
|  | HIS, B chain 368 | | D ring | pi-alkyl | 4.66 |
|  | PHE, A chain 242 | | C18 | pi-alkyl | 5.28 |
|  | PHE, B chain 366 | | B ring | pi-alkyl | 5.39 |
| **Alpha 1-antichymotrypsin variant NewBG-III (6hgk)** | ALA, chain A 274 | | C19 | alkyl | 3.83 |
|  | LEU, Chain A 273 | | B ring | alkyl | 5.25 |
|  | LEU, Chain A 273 | | C19 | alkyl | 4.25 |
|  | TRP, chain B 386 | | B ring | pi-alkyl | 4.5 |
|  | TRP, chain B 386 | | B ring | pi-alkyl | 4.67 |
|  | TRP, chain B 386 | | between C and D rings (C14) | pi-sigma | 3.88 |
|  | TRP, chain B 386 | | D ring | pi-alkyl | 4.83 |
|  | PHE, chain A 252 | | D ring | pi-alkyl | 5.41 |
|  | HIS, chain B 383 | | C_20_=O | carbon hydrogen bond | 3.4 |
|  | HIS, chain B 383 | | D ring | pi-alkyl | 4.71 |
|  | VAL, chain B 381 | | D ring | alkyl | 4.12 |
|  | VAL, chain A 31 | | B ring | alkyl | 5.15 |

**Table S16.** (B): Consensus binding residues of the pocket and interactions between progesterone and the steroid binding proteins.

| Involved amino acid residue | Parent protein(s) | Average Residue number per pocket | Involved P4 site(s) | Interaction type | Average Interaction distance (Å) |
| --- | --- | --- | --- | --- | --- |
| TRP (B chain) | both | 1 | A ring, B ring, C ring, D ring, between C and D rings (C14) | pi-alkyl, pi-sigma | 4.75 (for pi-alkyl bond), 3.79 (for pi-sigma bond) |
| HIS (B chain) | both | 1 | D ring, C20=O | pi-alkyl, carbon hydrogen bond | 4.68 (for pi-alkyl bond), 3.4 (for carbon hydrogen bond) |
| PHE (A chain and B chain) | both | 1.5 | C18, B ring, D ring | pi-alkyl | 5.36 |

**Table S17**. (A): Details of biochemical interactions between progesterone and cytochrome P450 enzymes.

| **Protein (PDB ID)** | **Involved amino acid residue (number)** | | **Involved P4 site** | **Interaction type** | **Interaction distance (Å)** |
| --- | --- | --- | --- | --- | --- |
| **Cytochrome P450 CYP260A1 (S276I) (6f8c)** | | VAL 163 | C19 | alkyl | 5.27 |
|  |  | LEU 159 | C19 | alkyl | 5.48 |
|  |  | LEU 162 | C19 | alkyl | 5.02 |
|  |  | LEU 162 | C18 | alkyl | 5.26 |
|  |  | LEU 228 | C18 | alkyl | 4.72 |
|  |  | LEU 69 | C18 | alkyl | 4.68 |
| **Cytochrome P450 CYP260A1 (S276N) (6f88)** | | HOH 788 | C_3_=O | water conventional hydrogen bond | 2.87 |
|  |  | LEU 162 | C19 | alkyl | 5.4 |
|  |  | LEU 69 | C19 | alkyl | 4.97 |
|  |  | LEU 159 | C19 | alkyl | 5.43 |
|  |  | LEU 228 | C19 | alkyl | 4.58 |
|  |  | HOH 613 | C_20_=O | water conventional hydrogen bond | 2.5 |
|  |  | ALA 74 | B ring | alkyl | 5.42 |
|  |  | SER 225 | C_3_=O | carbon hydrogen bond | 2.98 |
| **Cytochrome P450 3A4 (5a1r)** | | ASP 214 | C_20_=O | conventional hydrogen bond | 2.88 |
|  |  | PHE 213 | D ring | pi-alkyl | 4.82 |
|  |  | VAL 240 | D ring | alkyl | 4.65 |
|  |  | PHE 220 | C18 | pi-alkyl | 4.74 |
|  |  | PHE 220 | D ring | pi-alkyl | 4.79 |
|  |  | PHE 219 | C19 | pi-alkyl | 4.41 |
|  |  | PHE 219 | B ring | pi-alkyl | 4.52 |
|  |  | PHE 219 | C18 | pi-alkyl | 5.15 |
| **Cytochrome P450 3A4 (Cyp3A4 monooxygenase) (5a1p)** | | ASP 214 | C_20_=O | conventional hydrogen bond | 3.03 |
|  |  | PHE 213 | D ring | pi-alkyl | 4.84 |
|  |  | VAL 240 | D ring | alkyl | 4.9 |
|  |  | PHE 220 | D ring | pi-alkyl | 4.94 |
|  |  | PHE 219 | C19 | pi-alkyl | 4.72 |
|  |  | PHE 219 | B ring | pi-alkyl | 4.69 |
| **Human cytochrome P450 3A4 (1w0f)** | | PHE 219 | C19 | Pi-Alkyl | 4.53 |
|  |  | PHE 219 | B ring | Pi-Alkyl | 4.54 |
|  |  | PHE 219 | C18 | Pi-Alkyl | 4.79 |
|  |  | PHE 213 | D ring | Pi-Alkyl | 5.2 |
|  |  | PHE 213 | C_20_=O | Carbone hydrogen bond | 3.19 |
|  |  | PHE 220 | C18 | Pi-Alkyl | 4.66 |
|  |  | PHE 220 | D ring | Pi-Alkyl | 5.21 |
|  |  | ASP 214 | C_20_=O | conventional hydrogen bond | 2.96 |
|  |  | VAL 240 | D ring | Alkyl | 4.63 |
| **Human cytochrome P450 21A2 (4y8w)** | | ARG 234 | C_3_=O | conventional hydrogen bond | 2.73 |
|  |  | LEU 110 | C19 | alkyl | 4.69 |
|  |  | TRP 202 | C19 | pi-sigma | 3.68 |
|  |  | TRP 202 | C19 | pi-sigma | 3.9 |
|  |  | LEU 364 | C18 | alkyl | 4.19 |
|  |  | VAL 470 | C18 | alkyl | 5.15 |
|  |  | ILE 291 | C ring | alkyl | 5.47 |
|  |  | ILE 291 | B ring | alkyl | 4.69 |
| **Zebra fish cytochrome P450 17A2 (4r21)** | | ILE 219 | C19 | alkyl | 4.77 |
|  |  | VAL 480 | C19 | alkyl | 4.63 |
|  |  | SER 367 | C_20_=O | conventional hydrogen bond | 3.15 |
|  |  | ALA 302 | D ring | alkyl | 4.25 |
|  |  | ALA 302 | C ring | alkyl | 5.06 |
|  |  | ILE 371 | D ring | alkyl | 5.12 |
|  |  | ALA 120 | D ring | alkyl | 4.12 |
| **CYP154C5 from Nocardia farcinica (4j6c)** | | HOH 634 | C_3_=O | water conventional hydrogen bond | 3.3 |
|  |  | HOH 859 | C_3_=O | water conventional hydrogen bond | 3.02 |
|  |  | HOH 762 | C_3_=O | water conventional hydrogen bond | 2.7 |
|  |  | MET 84 | C19 | alkyl | 4.51 |
|  |  | HEM 502 | C21 | pi-sigma | 3.73 |
|  |  | HEM 502 | D ring | pi-alkyl | 3.87 |
|  |  | HEM 502 | D ring | pi-alkyl | 4.6 |
|  |  | GLN 398 | C_20_=O | conventional hydrogen bond | 2.9 |
|  |  | ALA 244 | D ring | alkyl | 3.76 |
|  |  | ALA 244 | C ring | alkyl | 4.76 |
|  |  | PHE 92 | D ring | pi-alkyl | 5.24 |
|  |  | PHE 92 | C19 | pi-alkyl | 4.89 |
|  |  | PHE 92 | B ring | pi-alkyl | 4.51 |
|  |  | ALA 243 | B ring | alkyl | 4.54 |
|  |  | ALA 240 | B ring | alkyl | 4.69 |

**Table S17**. (B): Consensus binding residues of the pocket and interactions between progesterone and the cytochrome P450 enzymes.

| Involved amino acid residue | Parent protein(s) | Average Residue number per pocket | Involved P4 site(s) | Interaction type | Average Interaction distance (Å) |
| --- | --- | --- | --- | --- | --- |
| VAL | all excluding CYP154C5 from Nocardia farcinica (4j6c) and Cytochrome P450 CYP260A1 (S276N) (6f88) | 0.75 | C18, C19, D ring | alkyl | 4.87 |
| LEU | human cytochrome P450 21A2 (4y8w), Cytochrome P450 CYP260A1 (S276N) (6f88), and Cytochrome P450 CYP260A1 (S276I) (6f8c) | 1.25 | C18, C19 | alkyl | 4.95 |
| ALA | cytochrome P450 CYP260A1 (S276N) (6f88), Zebra fish cytochrome P450 17A2 (4r21), CYP154C5 from Nocardia farcinica (4j6c) | 0.75 | B ring, C ring, D ring | alkyl | 4.58 |
| PHE | cytochrome P450 3A4 (5a1r), CYP154C5 from Nocardia farcinica (4j6c), Human cytochrome P450 3A4 (1w0f), Cytochrome P450 3A4 (5a1r) | 1.25 | C18, C19, B ring, D ring, C_20_=O | pi-alkyl, carbon hydrogen bond | 3.19 (carbon hydrogen bond), 4.8 (pi-alkyl) |

**Table S18**. (A): Details of biochemical interactions between progesterone and other P4-binding proteins.

| **Protein (PDB ID)** | | **Involved amino acid residue (number)** | | **Involved P4 site** | **Interaction type** | **Interaction distance (Å)** |
| --- | --- | --- | --- | --- | --- | --- |
| **Human apolipoprotein D (2hzq)** | ILE 70 | | | C19 | alkyl | 4.83 |
|  | PHE 89 | | | C19 | pi-sigma | 3.71 |
|  | PHE 89 | | | B ring | pi-alkyl | 4.89 |
|  | PHE 89 | | | C18 | pi-alkyl | 5.47 |
|  | ALA 44 | | | C18 | alkyl | 4.07 |
|  | TYR 46 | | | C_20_=O | conventional hydrogen bond | 2.58 |
|  | TYR 98 | | | D ring | pi-alkyl | 4.89 |
|  | VAL 56 | | | C18 | alkyl | 5.38 |
|  | TRP 127 | | | D ring | pi-alkyl | 4.29 |
|  | TRP 127 | | | D ring | pi-alkyl | 4.14 |
|  | TRP 127 | | | C ring | pi-alkyl | 4.54 |
|  | TRP 127 | | | C ring | pi-alkyl | 5.22 |
|  | TRP 127 | | | B ring | pi-alkyl | 5.07 |

| **Engineered avidin (5lur)** | PHE 72 | C19 | pi-alkyl | 4.86 |
| --- | --- | --- | --- | --- |
|  | ALA 35 | C19 | alkyl | 4.02 |
|  | TRP 70 | C19 | pi-alkyl | 4.12 |
|  | TRP 70 | B ring | pi-alkyl | 4.55 |
|  | TRP 70 | B ring | pi-alkyl | 5.02 |
|  | TRP 70 | C18 | pi-alkyl | 5.2 |
|  | HIS 16 | C ring | pi-alkyl | 5.27 |
|  | HIS 16 | C18 | pi-alkyl | 4.54 |
|  | TYR 33 | C18 | pi-alkyl | 4.91 |
|  | ASN 118 | C_20_=O | conventional hydrogen bond | 2.44 |
|  | TRP 97 | C_20_=O | conventional hydrogen bond | 2.98 |
|  | TRP 97 | D ring | pi-alkyl | 4.22 |
|  | TRP 97 | D ring | pi-alkyl | 4.56 |
|  | LEU 99 | D ring | alkyl | 4.62 |

**Table S18.** (B): Consensus binding residues of the pocket and interactions in other P4-binding proteins.

| Involved amino acid residue | Parent protein(s) | Average Residue number per pocket | Involved P4 site(s) | Interaction type | Average Interaction distance (Å) |
| --- | --- | --- | --- | --- | --- |
| PHE | Human apolipoprotein D (2hzq), Engineered avidin (5lur) | 1 | C18, C19, B ring | pi-sigma, pi-alkyl | 3.71 (for pi-sigma bond), 5.07 (for pi-alkyl bond) |
| ALA | Human apolipoprotein D (2hzq), Engineered avidin (5lur) | 1 | C18, C19 | alkyl | 4.045 |
| TYR | Human apolipoprotein D (2hzq), Engineered avidin (5lur) | 1.5 | C18, C20=O, D ring | conventional hydrogen bond, pi-alkyl | 2.58 (for conventional hydrogen bond), 4.9 (for pi-alkyl bond) |
| TRP | Human apolipoprotein D (2hzq), Engineered avidin (5lur) | 1.5 | C18, C19, B ring, C ring, D ring, C20=O | conventional hydrogen bond, pi-alkyl | 2.98 (for conventional hydrogen bond), 4.63 (for pi-alkyl bond) |

**Table S19.** Critical residues of P4-binding proteins pockets predicted by the in silico alanine scanning mutagenesis.

| Protein | Pocket Residues | delta-AutoDock4.1Score |
| --- | --- | --- |
| human progesterone receptor ligand binding domain (1a28) | GLN_725_ALA | -0.7507 |
|  | PHE_778_ALA | -0.6457 |
|  | MET_756_ALA | -0.6452 |
|  | LEU_718_ALA | -0.4707 |
|  | TYR_890_ALA | -0.468 |
|  | TRP_755_ALA | -0.3762 |
|  | MET_759_ALA | -0.3516 |
|  | MET_801_ALA | -0.3495 |
|  | MET_909_ALA | -0.3482 |
|  | ASN_719_ALA | -0.3422 |
|  | LEU_887_ALA | -0.3326 |
|  | PHE_905_ALA | -0.3263 |
|  | LEU_715_ALA | -0.3155 |
|  | LEU_763_ALA | -0.3065 |
|  | LEU_721_ALA | -0.302 |
|  | LEU_797_ALA | -0.2769 |
|  | ARG_766_ALA | -0.1848 |
|  | CYS_891_ALA | -0.1634 |
|  | VAL_760_ALA | -0.1408 |
|  | THR_894_ALA | -0.1036 |
|  | VAL_903_ALA | -0.0606 |
|  | GLY_722_ALA | 0.2761 |

| Protein | Pocket Residues | delta-AutoDock4.1Score |
| --- | --- | --- |
| anti-progesterone antibody DB3 (complex with P4) (1dbb) | GLY_33_ALA | -1.555 |
|  | SER_92_ALA | -1.5078 |
|  | SER_91_ALA | -0.8512 |
|  | PRO_96_ALA | -0.4466 |
|  | TYR_100A_ALA | -0.3894 |
|  | TRP_50_ALA | -0.3367 |
|  | PHE_100B_ALA | -0.2824 |
|  | GLY_95_ALA | -0.2559 |
|  | TRP_100_ALA | -0.0918 |
|  | ASN_35_ALA | -0.036 |
|  | VAL_94_ALA | -0.0237 |
|  | ASP_96_ALA | 0.0061 |
|  | TYR_97_ALA | 0.0483 |
|  | HIS_93_ALA | 0.0523 |
|  | HIS_27D_ALA | 0.0962 |

| Protein | Pocket Residues | delta-AutoDock4.1Score |
| --- | --- | --- |
| anti-progesterone antibody DB3 (complex with 11-hemmisuccinate-P4) (1dbm) | TRP_100_ALA | -2.842 |
|  | TRP_50_ALA | -1.8679 |
|  | TYR_97_ALA | -0.7554 |
|  | ASN_35_ALA | -0.5372 |
|  | PHE_100B_ALA | -0.415 |
|  | VAL_94_ALA | -0.3462 |
|  | PRO_96_ALA | -0.3225 |
|  | HIS_27D_ALA | -0.2573 |
|  | GLY_95_ALA | -0.1986 |
|  | ASN_52_ALA | -0.0419 |
|  | SER_92_ALA | -0.0281 |
|  | TYR_32_ALA | -0.0122 |
|  | ASP_96_ALA | 0.0331 |
|  | GLY_33_ALA | 0.1151 |
|  | SER_91_ALA | 0.1442 |

| Protein | Pocket Residues | delta-AutoDock4.1Score |
| --- | --- | --- |
| pentaerythritol tetranitrate reductase (PETN reductase) (1h60) | TYR_351_ALA | -1.309 |
|  | TYR_68_ALA | -0.923 |
|  | TYR_186_ALA | -0.784 |
|  | HIS_184_ALA | -0.7232 |
|  | GLN_241_ALA | -0.5636 |
|  | HIS_181_ALA | -0.4941 |
|  | ARG_142_ALA | -0.329 |
|  | TRP_102_ALA | -0.3219 |
|  | LEU_275_ALA | -0.3098 |
|  | THR_26_ALA | -0.1848 |
|  | ARG_130_ALA | -0.0093 |
|  | SER_132_ALA | 0.0316 |

| Protein | Pocket Residues | delta-AutoDock4.1Score |
| --- | --- | --- |
| human 20alpha-hydroxysteroid dehydrogenase (1mrq) | TRP_227_ALA | -1.8764 |
|  | TYR_24_ALA | -1.3231 |
|  | HIS_222_ALA | -1.0183 |
|  | LEU_54_ALA | -0.9895 |
|  | TYR_55_ALA | -0.5113 |
|  | ILE_129_ALA | -0.5079 |
|  | LEU_306_ALA | -0.4182 |
|  | LEU_308_ALA | -0.2777 |
|  | VAL_128_ALA | -0.1957 |
|  | ILE_310_ALA | -0.148 |
|  | GLU_224_ALA | -0.1108 |
|  | GLU_127_ALA | 0.0539 |

| Protein | Pocket Residues | delta-AutoDock4.1Score |
| --- | --- | --- |
| human cytochrome P450 3A4 (1w0f) | PHE_219_ALA | -1.3909 |
|  | PHE_220_ALA | -0.7055 |
|  | PHE_213_ALA | -0.4994 |
|  | ASP_217_ALA | -0.2707 |
|  | VAL_240_ALA | -0.2338 |
|  | ILE_238_ALA | -0.1826 |
|  | ARG_212_ALA | -0.0158 |
|  | ASP_214_ALA | 0.1324 |

| Protein | Pocket Residues | delta-AutoDock4.1Score |
| --- | --- | --- |
| mineralocorticoid receptor (2aa5) | PHE_829_ALA | -0.7881 |
|  | MET_807_ALA | -0.6782 |
|  | MET_845_ALA | -0.5815 |
|  | PHE_941_ALA | -0.5405 |
|  | LEU_769_ALA | -0.4583 |
|  | TRP_806_ALA | -0.4495 |
|  | GLN_776_ALA | -0.4194 |
|  | ARG_817_ALA | -0.3919 |
|  | ASN_770_ALA | -0.3625 |
|  | MET_852_ALA | -0.361 |
|  | LEU_814_ALA | -0.356 |
|  | LEU_938_ALA | -0.3243 |
|  | LEU_766_ALA | -0.3061 |
|  | PHE_956_ALA | -0.2605 |
|  | LEU_772_ALA | -0.2397 |
|  | CYS_942_ALA | -0.1696 |
|  | THR_945_ALA | -0.086 |
|  | VAL_954_ALA | -0.0663 |
|  | ALA_773_ALA | 0.0023 |
|  | SER_811_ALA | 0.0074 |
|  | SER_810_ALA | 0.0319 |

| Protein | Pocket Residues | delta-AutoDock4.1Score |
| --- | --- | --- |
| pentaerythritol tetranitrate reductase (2aba) | TYR_351_ALA | -1.2981 |
|  | ARG_142_ALA | -1.0733 |
|  | TYR_68_ALA | -1.0533 |
|  | TYR_186_ALA | -0.8139 |
|  | HIS_184_ALA | -0.7094 |
|  | GLN_241_ALA | -0.575 |
|  | HIS_181_ALA | -0.5269 |
|  | TRP_102_ALA | -0.3419 |
|  | LEU_275_ALA | -0.2311 |
|  | THR_26_ALA | -0.1915 |
|  | ARG_130_ALA | -0.0559 |
|  | THR_131_ALA | -0.0412 |
|  | SER_132_ALA | 0.0282 |

| Protein | Pocket Residues | delta-AutoDock4.1Score |
| --- | --- | --- |
| mineralocorticoid receptor S810L mutant (2aa6) | PHE_829_ALA | -0.8022 |
|  | MET_807_ALA | -0.6845 |
|  | MET_845_ALA | -0.6435 |
|  | PHE_941_ALA | -0.6092 |
|  | LEU_769_ALA | -0.4894 |
|  | GLN_776_ALA | -0.4263 |
|  | ASN_770_ALA | -0.3727 |
|  | LEU_814_ALA | -0.3709 |
|  | TRP_806_ALA | -0.3599 |
|  | LEU_938_ALA | -0.3564 |
|  | MET_852_ALA | -0.3482 |
|  | LEU_766_ALA | -0.3278 |
|  | LEU_810_ALA | -0.3206 |
|  | PHE_956_ALA | -0.3161 |
|  | ARG_817_ALA | -0.3137 |
|  | LEU_772_ALA | -0.2362 |
|  | THR_945_ALA | -0.1911 |
|  | CYS_942_ALA | -0.1503 |
|  | VAL_954_ALA | -0.0704 |
|  | ALA_773_ALA | 0.008 |
|  | SER_811_ALA | 0.008 |

| Protein | Pocket Residues | delta-AutoDock4.1Score |
| --- | --- | --- |
| delta(4)-3-ketosteroid 5beta-reductase (akr1d1) (3cot) | TRP_230_ALA | -1.6867 |
|  | TYR_58_ALA | -1.1222 |
|  | TYR_132_ALA | -1.0279 |
|  | TYR_26_ALA | -0.939 |
|  | TRP_314_ALA | -0.5502 |
|  | TRP_89_ALA | -0.4558 |
|  | LEU_311_ALA | -0.3875 |
|  | MET_313_ALA | -0.242 |
|  | TRP_140_ALA | -0.1666 |
|  | LYS_87_ALA | -0.162 |
|  | GLU_120_ALA | -0.1487 |
|  | ILE_229_ALA | -0.1251 |
|  | VAL_309_ALA | -0.0936 |

| Protein | Pocket Residues | delta-AutoDock4.1Score |
| --- | --- | --- |
| human apolipoprotein D (ApoD) (2hzq) | TRP_127_ALA | -1.9822 |
|  | PHE_89_ALA | -1.1576 |
|  | TYR_98_ALA | -0.4682 |
|  | LEU_129_ALA | -0.4574 |
|  | TYR_46_ALA | -0.384 |
|  | ASN_58_ALA | -0.3775 |
|  | VAL_87_ALA | -0.2124 |
|  | ILE_70_ALA | -0.2037 |
|  | THR_34_ALA | -0.1957 |
|  | GLU_28_ALA | -0.1487 |
|  | ILE_42_ALA | -0.1123 |
|  | SER_132_ALA | 0.0282 |

| Protein | Pocket Residues | delta-AutoDock4.1Score |
| --- | --- | --- |
| corticosteroid-binding globulin (4bb2) | TRP_371_ALA | -2.6206 |
|  | PHE_366_ALA | -0.7685 |
|  | GLN_232_ALA | -0.7657 |
|  | HIS_368_ALA | -0.5491 |
|  | ILE_263_ALA | -0.525 |
|  | ARG_260_ALA | -0.5046 |
|  | PHE_242_ALA | -0.4365 |
|  | ASN_264_ALA | -0.4297 |
|  | VAL_22_ALA | -0.2555 |
|  | THR_240_ALA | -0.1108 |
|  | SER_267_ALA | -0.0559 |
|  | SER_19_ALA | -0.0068 |
|  | ALA_18_ALA | 0.0199 |

| Protein | Pocket Residues | delta-AutoDock4.1Score |
| --- | --- | --- |
| 1E9 LeuH47Trp/ArgH100Trp anti-P4 antibody (2o5y) | TRP_50_ALA | -1.897 |
|  | PRO_96_ALA | -1.6209 |
|  | SER_91_ALA | -0.4425 |
|  | PHE_89_ALA | -0.3904 |
|  | MET_100B_ALA | -0.305 |
|  | TRP_100_ALA | -0.2648 |
|  | THR_97_ALA | -0.2536 |
|  | TRP_47_ALA | -0.1363 |
|  | ALA_100A_ALA | -0.0486 |
|  | GLY_95_ALA | 0.0122 |
|  | ASN_35_ALA | 0.1118 |

| Protein | Pocket Residues | delta-AutoDock4.1Score |
| --- | --- | --- |
| ancestral 3-keto steroid receptor (4fn9) | GLN_41_ALA | -0.7629 |
|  | PHE_94_ALA | -0.5347 |
|  | LEU_34_ALA | -0.3988 |
|  | ASN_35_ALA | -0.3904 |
|  | MET_75_ALA | -0.3468 |
|  | TRP_71_ALA | -0.3322 |
|  | LEU_37_ALA | -0.2803 |
|  | PHE_206_ALA | -0.2623 |
|  | ARG_82_ALA | -0.2515 |
|  | PHE_221_ALA | -0.2277 |
|  | MET_72_ALA | -0.2052 |
|  | MET_110_ALA | -0.1355 |
|  | THR_210_ALA | -0.095 |
|  | LEU_31_ALA | -0.0231 |
|  | ALA_76_ALA | -0.0181 |
|  | LEU_203_ALA | -0.0167 |
|  | ALA_38_ALA | -0.0086 |
|  | MET_79_ALA | 0.0359 |
|  | MET_117_ALA | 0.0427 |
|  | CYS_207_ALA | 0.1781 |

| Protein | Pocket Residues | delta-AutoDock4.1Score |
| --- | --- | --- |
| CYP154C5 from *Nocardia farcinica* (4j6c) | PHE_92_ALA | -1.4981 |
|  | GLN_398_ALA | -0.9095 |
|  | PHE_179_ALA | -0.6241 |
|  | PHE_180_ALA | -0.4228 |
|  | MET_84_ALA | -0.3971 |
|  | LEU_294_ALA | -0.3126 |
|  | VAL_87_ALA | -0.2964 |
|  | VAL_291_ALA | -0.1693 |
|  | THR_248_ALA | -0.1468 |
|  | GLN_239_ALA | -0.0709 |
|  | ALA_243_ALA | -0.0087 |
|  | GLY_83_ALA | -0.0045 |
|  | ALA_244_ALA | 0.0122 |
|  | ALA_240_ALA | 0.0161 |

| Protein | Pocket Residues | delta-AutoDock4.1Score |
| --- | --- | --- |
| Human 3-alpha Hydroxysteroid Dehydrogenase Type 3(4l1w) | TRP_227_ALA | -1.8471 |
|  | TYR_24_ALA | -1.1237 |
|  | TRP_86_ALA | -0.4737 |
|  | ILE_129_ALA | -0.4732 |
|  | VAL_54_ALA | -0.4261 |
|  | LEU_308_ALA | -0.3824 |
|  | TYR_55_ALA | -0.3415 |
|  | LEU_306_ALA | -0.2219 |
|  | VAL_128_ALA | -0.1865 |
|  | HIS_222_ALA | -0.1648 |
|  | GLU_224_ALA | -0.1568 |
|  | HIS_117_ALA | -0.1537 |

| Protein | Pocket Residues | delta-AutoDock4.1Score |
| --- | --- | --- |
| human 3-alpha hydroxysteroid dehydrogenase type 3 V54L mutant (4l1x) | TRP_227_ALA | -1.9243 |
|  | TYR_24_ALA | -1.2183 |
|  | LEU_54_ALA | -0.9222 |
|  | HIS_222_ALA | -0.8367 |
|  | ILE_129_ALA | -0.5231 |
|  | TYR_55_ALA | -0.4468 |
|  | LEU_306_ALA | -0.3453 |
|  | LEU_308_ALA | -0.3108 |
|  | VAL_128_ALA | -0.2277 |
|  | GLU_224_ALA | -0.0595 |

| Protein | Pocket Residues | delta-AutoDock4.1Score |
| --- | --- | --- |
| ancestral ketosteroid receptor (4ltw) | MET_72_ALA | -0.6847 |
|  | GLN_41_ALA | -0.6352 |
|  | PHE_94_ALA | -0.6253 |
|  | ASN_35_ALA | -0.3672 |
|  | MET_75_ALA | -0.3394 |
|  | LEU_34_ALA | -0.3357 |
|  | PHE_206_ALA | -0.2826 |
|  | LEU_37_ALA | -0.2747 |
|  | MET_110_ALA | -0.18 |
|  | MET_79_ALA | -0.163 |
|  | LEU_203_ALA | -0.1582 |
|  | TRP_71_ALA | -0.0848 |
|  | THR_210_ALA | -0.0698 |
|  | MET_117_ALA | -0.053 |
|  | LEU_31_ALA | -0.0471 |
|  | ARG_82_ALA | -0.0391 |
|  | ALA_76_ALA | -0.0121 |
|  | ALA_38_ALA | 0.0019 |
|  | CYS_207_ALA | 0.0692 |

| Protein | Pocket Residues | delta-AutoDock4.1Score |
| --- | --- | --- |
| human steroidogenic cytochrome P450 17A1 mutant A105L (complex with P4) (4nkx) | GLY_301_ALA | -0.8459 |
|  | PHE_114_ALA | -0.7982 |
|  | LEU_105_ALA | -0.5788 |
|  | ILE_205_ALA | -0.3609 |
|  | ILE_206_ALA | -0.3439 |
|  | VAL_482_ALA | -0.3407 |
|  | ILE_371_ALA | -0.3261 |
|  | VAL_483_ALA | -0.2689 |
|  | ASN_202_ALA | -0.2036 |
|  | LEU_209_ALA | -0.1862 |
|  | GLU_305_ALA | -0.1779 |
|  | THR_306_ALA | -0.1612 |
|  | ASP_298_ALA | -0.143 |
|  | VAL_366_ALA | -0.0945 |
|  | ALA_113_ALA | -0.022 |
|  | ALA_302_ALA | -0.0138 |
|  | GLY_297_ALA | 0.0329 |

| Protein | Pocket Residues | delta-AutoDock4.1Score |
| --- | --- | --- |
| human steroidogenic cytochrome P450 17A1 mutant A105L (complex with 17α-OH-progesterone) (4nky) | GLY_301_ALA | -0.9772 |
|  | PHE_114_ALA | -0.8261 |
|  | LEU_105_ALA | -0.5209 |
|  | VAL_482_ALA | -0.4214 |
|  | ILE_371_ALA | -0.4075 |
|  | ILE_206_ALA | -0.3927 |
|  | ILE_205_ALA | -0.3905 |
|  | VAL_483_ALA | -0.2427 |
|  | THR_306_ALA | -0.1743 |
|  | ASN_202_ALA | -0.1552 |
|  | GLU_305_ALA | -0.1432 |
|  | LEU_209_ALA | -0.1352 |
|  | ASP_298_ALA | -0.1111 |
|  | VAL_366_ALA | -0.1026 |
|  | ARG_239_ALA | -0.0608 |
|  | ALA_302_ALA | -0.0161 |
|  | ALA_367_ALA | -0.0139 |
|  | ALA_113_ALA | -0.0067 |
|  | GLY_297_ALA | 0.0193 |

| Protein | Pocket Residues | delta-AutoDock4.1Score |
| --- | --- | --- |
| human steroidogenic cytochrome P450 17A1 mutant A105L (complex with 17α-OH-progesterone) (4nkz) | PHE_114_ALA | -0.8748 |
|  | LEU_105_ALA | -0.4452 |
|  | ILE_371_ALA | -0.4334 |
|  | VAL_482_ALA | -0.4158 |
|  | ILE_206_ALA | -0.3491 |
|  | ILE_205_ALA | -0.3251 |
|  | VAL_483_ALA | -0.2766 |
|  | GLY_301_ALA | -0.2629 |
|  | THR_306_ALA | -0.2213 |
|  | ASN_202_ALA | -0.1514 |
|  | LEU_209_ALA | -0.1443 |
|  | VAL_366_ALA | -0.1089 |
|  | ASP_298_ALA | -0.0736 |
|  | ALA_367_ALA | -0.0094 |
|  | ALA_113_ALA | -0.0071 |
|  | GLY_297_ALA | 0.0127 |
|  | ALA_302_ALA | 0.0163 |

| Protein | Pocket Residues | delta-AutoDock4.1Score |
| --- | --- | --- |
| zebra fish cytochrome P450 17A2 (4r21) | PHE_121_ALA | -0.8182 |
|  | ILE_371_ALA | -0.4966 |
|  | VAL_480_ALA | -0.3888 |
|  | ILE_216_ALA | -0.3038 |
|  | ILE_212_ALA | -0.225 |
|  | GLU_298_ALA | -0.201 |
|  | THR_306_ALA | -0.132 |
|  | VAL_213_ALA | -0.1201 |
|  | VAL_366_ALA | -0.0585 |
|  | SER_367_ALA | -0.053 |
|  | ALA_120_ALA | -0.0251 |
|  | ALA_302_ALA | 0.0124 |
|  | GLY_301_ALA | 0.083 |

| Protein | Pocket Residues | delta-AutoDock4.1Score |
| --- | --- | --- |
| cytochrome P450 3A4 (5a1p) | PHE_219_ALA | -1.3304 |
|  | PHE_220_ALA | -0.8032 |
|  | PHE_213_ALA | -0.4919 |
|  | ASP_217_ALA | -0.3535 |
|  | ILE_238_ALA | -0.1843 |
|  | VAL_240_ALA | -0.1769 |
|  | ASP_214_ALA | -0.058 |
|  | ARG_212_ALA | -0.0037 |
|  | CYS_239_ALA | 0.0129 |

| Protein | Pocket Residues | delta-AutoDock4.1Score |
| --- | --- | --- |
| human cytochrome P450 21A2 (4y8w) | TRP_202_ALA | -1.5235 |
|  | ILE_291_ALA | -0.6869 |
|  | ARG_234_ALA | -0.4909 |
|  | LEU_110_ALA | -0.4774 |
|  | VAL_101_ALA | -0.4642 |
|  | LEU_364_ALA | -0.4536 |
|  | LEU_199_ALA | -0.3898 |
|  | VAL_198_ALA | -0.235 |
|  | VAL_360_ALA | -0.2266 |
|  | VAL_470_ALA | -0.209 |
|  | VAL_287_ALA | -0.2023 |
|  | ASP_288_ALA | -0.1644 |
|  | ILE_231_ALA | -0.1613 |
|  | ASP_107_ALA | -0.114 |
|  | ILE_471_ALA | -0.1116 |
|  | SER_109_ALA | -0.1102 |
|  | THR_296_ALA | -0.1051 |
|  | GLY_292_ALA | 0.1536 |

| Protein | Pocket Residues | delta-AutoDock4.1Score |
| --- | --- | --- |
| cytochrome P450 3A4 (5a1r) | PHE_219_ALA | -1.3678 |
|  | PHE_220_ALA | -0.7875 |
|  | PHE_213_ALA | -0.4675 |
|  | ASP_217_ALA | -0.2294 |
|  | VAL_240_ALA | -0.2073 |
|  | ILE_238_ALA | -0.1741 |
|  | CYS_239_ALA | -0.0023 |
|  | ARG_212_ALA | -0.0001 |
|  | ASP_214_ALA | 0.0289 |

| Protein | Pocket Residues | delta-AutoDock4.1Score |
| --- | --- | --- |
| plantago major multifunctional oxidoreductase mutant V150M (5mlm) | LYS_147_ALA | -1.0696 |
|  | ASN_205_ALA | -0.8054 |
|  | MET_150_ALA | -0.6406 |
|  | VAL_347_ALA | -0.5649 |
|  | PHE_153_ALA | -0.5344 |
|  | PHE_343_ALA | -0.4863 |
|  | ILE_156_ALA | -0.4378 |
|  | ILE_350_ALA | -0.3995 |
|  | PRO_353_ALA | -0.3567 |
|  | MET_215_ALA | -0.2108 |
|  | ARG_146_ALA | -0.2106 |
|  | CYS_352_ALA | -0.1781 |
|  | TRP_284_ALA | -0.1403 |
|  | PRO_351_ALA | -0.1162 |
|  | SER_248_ALA | -0.0142 |
|  | ALA_243_ALA | -0.0018 |
|  | ALA_346_ALA | 0.0068 |

| Protein | Pocket Residues | delta-AutoDock4.1Score |
| --- | --- | --- |
| human cytochrome P450 21A2 (complex with 17α-OH-progesterone) (5vbu) | TRP_202_ALA | -1.3442 |
|  | ILE_291_ALA | -0.6858 |
|  | LEU_364_ALA | -0.5029 |
|  | LEU_110_ALA | -0.4723 |
|  | VAL_101_ALA | -0.4623 |
|  | ARG_234_ALA | -0.4563 |
|  | LEU_199_ALA | -0.3345 |
|  | VAL_198_ALA | -0.2305 |
|  | VAL_360_ALA | -0.2101 |
|  | VAL_287_ALA | -0.1984 |
|  | VAL_470_ALA | -0.1791 |
|  | ASP_288_ALA | -0.1721 |
|  | ILE_231_ALA | -0.1689 |
|  | ASP_107_ALA | -0.1206 |
|  | SER_109_ALA | -0.1132 |
|  | THR_296_ALA | -0.0843 |
|  | GLY_292_ALA | 0.2165 |

| Protein | Pocket Residues | delta-AutoDock4.1Score |
| --- | --- | --- |
| cytochrome P450 CYP260A1 mutant S276I (6f8c) | LEU_69_ALA | -0.5968 |
|  | PHE_277_ALA | -0.5481 |
|  | LEU_159_ALA | -0.4982 |
|  | PHE_64_ALA | -0.459 |
|  | LEU_228_ALA | -0.3412 |
|  | LEU_162_ALA | -0.2918 |
|  | ILE_276_ALA | -0.2423 |
|  | THR_233_ALA | -0.2195 |
|  | VAL_163_ALA | -0.1871 |
|  | VAL_382_ALA | -0.1545 |
|  | VAL_279_ALA | -0.1188 |
|  | ALA_74_ALA | -0.001 |
|  | SER_225_ALA | 0.006 |
|  | GLY_278_ALA | 0.112 |
|  | GLY_229_ALA | 0.1446 |

| Protein | Pocket Residues | delta-AutoDock4.1Score |
| --- | --- | --- |
| P450 CYP260A1 mutant S276N (6f88) | LEU_69_ALA | -0.5277 |
|  | PHE_64_ALA | -0.4966 |
|  | PHE_277_ALA | -0.4118 |
|  | LEU_159_ALA | -0.4053 |
|  | VAL_382_ALA | -0.3286 |
|  | ASN_276_ALA | -0.285 |
|  | LEU_228_ALA | -0.2589 |
|  | VAL_163_ALA | -0.2533 |
|  | LEU_162_ALA | -0.2384 |
|  | THR_233_ALA | -0.2166 |
|  | LEU_280_ALA | -0.1809 |
|  | TYR_376_ALA | -0.0877 |
|  | SER_225_ALA | -0.0124 |
|  | ALA_74_ALA | -0.0058 |
|  | GLY_278_ALA | 0.0641 |
|  | GLY_229_ALA | 0.1877 |

| Protein | Pocket Residues | delta-AutoDock4.1Score |
| --- | --- | --- |
| plantago major multifunctional oxidoreductase (6gsd) | ASN_205_ALA | -0.7721 |
|  | LYS_147_ALA | -0.6539 |
|  | ILE_350_ALA | -0.5604 |
|  | PHE_153_ALA | -0.5262 |
|  | ARG_146_ALA | -0.4584 |
|  | PHE_343_ALA | -0.3594 |
|  | ILE_156_ALA | -0.3419 |
|  | PRO_353_ALA | -0.3231 |
|  | CYS_352_ALA | -0.2523 |
|  | VAL_150_ALA | -0.2462 |
|  | MET_215_ALA | -0.2427 |
|  | VAL_347_ALA | -0.1484 |
|  | TRP_284_ALA | -0.1255 |
|  | TYR_179_ALA | -0.086 |
|  | PRO_351_ALA | -0.0566 |
|  | ALA_346_ALA | -0.0174 |
|  | SER_248_ALA | -0.0082 |

| Protein | Pocket Residues | delta-AutoDock4.1Score | |
| --- | --- | --- | --- |
| alpha1-antichymotrypsin variant NewBG-III (6hgk) | TRP_386_ALA | -2.4668 | |
|  | PHE_252_ALA | | -0.7307 |
|  | LEU_273_ALA | | -0.5912 |
|  | ARG_270_ALA | | -0.4295 |
|  | HIS_383_ALA | | -0.3965 |
|  | VAL_31_ALA | | -0.3241 |
|  | VAL_381_ALA | | -0.2868 |
|  | GLN_242_ALA | | -0.0558 |
|  | ARG_24_ALA | | -0.0187 |
|  | ALA_274_ALA | | -0.0071 |
|  | SER_28_ALA | | 0.0007 |
|  | ALA_27_ALA | | 0.0212 |
|  | GLY_277_ALA | | 0.1504 |
